# Structure of the teneurin-latrophilin complex: Alternative splicing controls synapse specificity by a novel mechanism

**DOI:** 10.1101/2020.01.13.901090

**Authors:** Jingxian Li, Yuan Xie, Shaleeka Cornelius, Xian Jiang, Szymon P. Kordon, Richard Sando, Man Pan, Katherine Leon, Thomas C. Südhof, Minglei Zhao, Demet Araç

## Abstract

The trans-synaptic interaction of the cell-adhesion molecules teneurins (Tenm’s) with latrophilins (Lphn’s) promotes excitatory synapse formation when Lphn’s simultaneously interact with FLRTs. Insertion of a short alternatively-spliced region within Tenm’s abolishes the Tenm-Lphn interaction and switches Tenm function to specify inhibitory synapses. How Tenm’s bind to Lphn’s in a manner regulated by alternative splicing remains unclear. Here, we report the high-resolution cryo-EM structure of the Tenm2-Lphn3 complex, and describe the trimeric Tenm2-Lphn3-FLRT3 complex. The structure reveals that the N-terminal lectin-like domain of Lphn3 binds to the Tenm2 barrel at a site far away from the alternatively-spliced region. Alternative-splicing regulates the Tenm2-Lphn3 interaction by hindering access to the Lphn-binding surface rather than altering it. Strikingly, mutagenesis of the Lphn-binding surface of Tenm2 abolishes the Lphn3 interaction and impairs excitatory but not inhibitory synapse formation. These results suggest that a multi-level coincident binding mechanism mediated by a cryptic adhesion complex between Tenm’s and Lphn’s regulates synapse specificity.

## INTRODUCTION

Neural circuit assembly and function in the central nervous system requires precise formation and specification of diverse excitatory and inhibitory synapse subtypes. Imbalances in the ratio of excitatory to inhibitory synapse function are thought to be a major component of brain disorders such as autism, mental retardation and attention deficit hyperactivity disorder (Garber, 2007). Recent work suggested that combinatorial sets of trans-synaptic interactions between cell-adhesion molecular, including teneurins (Tenm’s) and latrophilins (Lphn’s), mediate synapse formation and regulate the exquisite specification of synapses, but the underlying molecular mechanisms remain largely unexplored (Li et al., 2018; Sando et al., 2019).

Tenm’s and Lphn’s are evolutionarily conserved cell-surface proteins. While the roles of Tenm’s and Lphn’s in early organisms remain unclear, they have critical roles in embryonic development and brain wiring in higher eukaryotes. Tenm’s (Tenm1-4 in mammals) are large type-II transmembrane proteins that are composed of an N-terminal cytoplasmic sequence, a single transmembrane region, and a large extracellular region (ECR) composed of >2000 amino acids with partial homology to bacterial Tc toxins (Fig. 1A) (Baumgartner and Chiquet-Ehrismann, 1993; Baumgartner et al., 1994; Levine et al., 1994; Oohashi et al., 1999). They form constitutive *cis*-dimers via highly conserved disulfide bonds formed in proximity to their transmembrane helix (Fig. 1C) (Feng et al., 2002; Oohashi et al., 1999; Vysokov et al., 2016). Tenm’s have central roles in tissue polarity, embryogenesis, heart development, axon guidance, and synapse formation (Berns et al., 2018; Boucard et al., 2014; Doyle et al., 2006; Hong et al., 2012; Levine et al., 1994; Mosca et al., 2012; Mosca and Luo, 2014; Silva et al., 2011; Woelfle et al., 2016); and are linked to various diseases including neurological disorders, developmental problems, various cancers and congenital general anosmia (Aldahmesh et al., 2012; Alkelai et al., 2016; Chassaing et al., 2016; Graumann et al., 2017; Hor et al., 2015; Talamillo et al., 2017; Ziegler et al., 2012). Lphn’s (Lphn1-3 in mammals) belong to the adhesion-type G-protein coupled receptor (GPCR) family (Krasnoperov et al., 1996; Lelianova et al., 1997) (Sugita et al., 1999) and have a large N-terminal ECR (>800 amino acids) in addition to their signaling seven-pass transmembrane domain and cytoplasmic tail (Fig. 1B) (Deak et al., 2009; Krasnoperov et al., 1997; Lelianova et al., 1997; Sudhof, 2001; Sugita et al., 1999). Lphn’s have roles in embryogenesis, tissue polarity, synaptic development and neural circuit connectivity, interestingly almost identical to the functions of Tenm’s (Anderson et al., 2017; Boucard et al., 2014; Deak et al., 2009; Hamann et al., 2015; Krasnoperov et al., 1997; Lelianova et al., 1997; Silva et al., 2011; Sudhof, 2001; Sugita et al., 1999) (Fig. 1B). Lphn3 mutations are linked to attention deficit hyperactivity disorder (ADHD), as well as numerous cancers in humans (Arcos-Burgos et al., 2010; Kan et al., 2010; O’Hayre et al., 2013).

**Figure 1:**
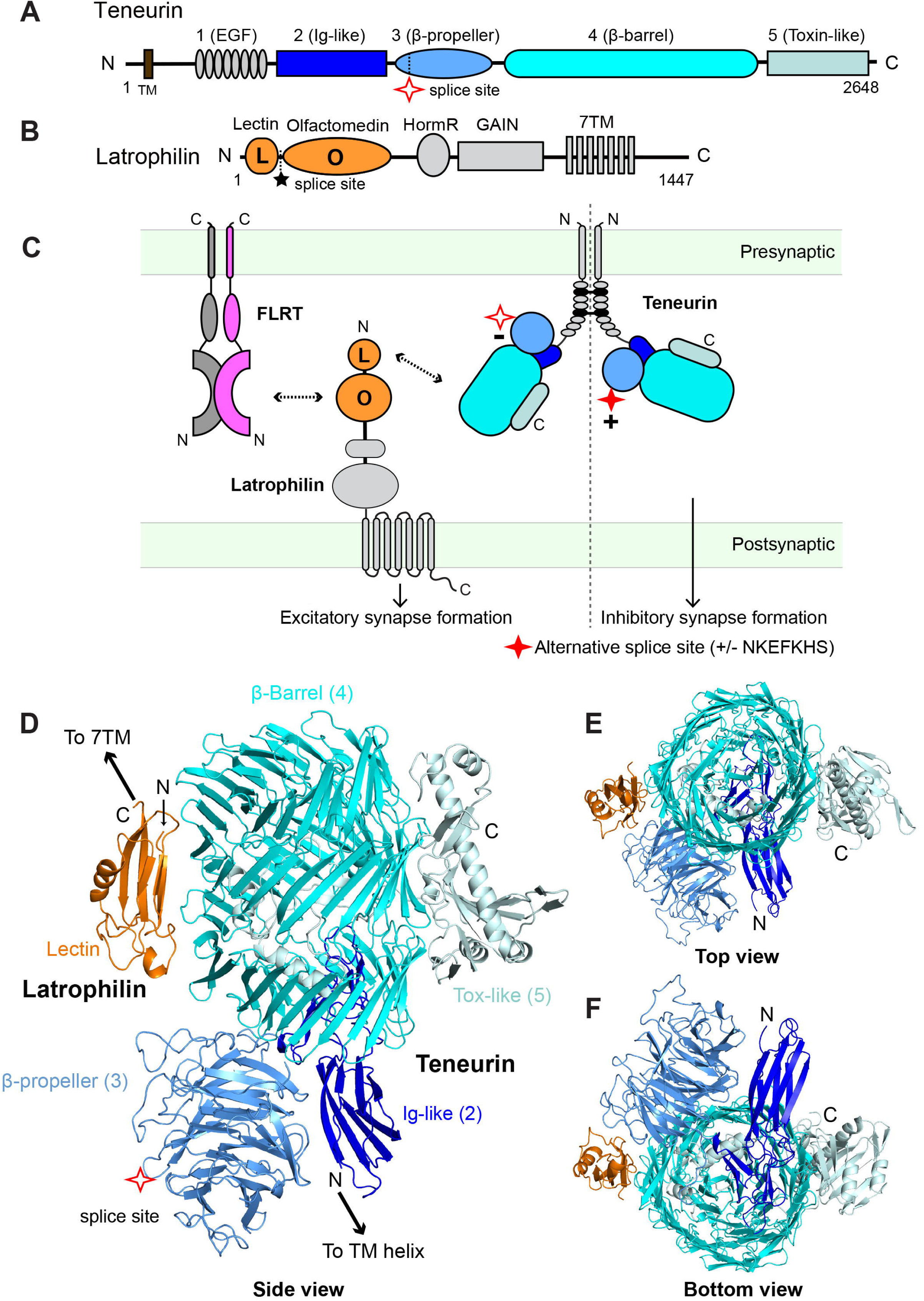
The structure of the Tenm2/Lphn3 complex. (A) Schematic diagram of human Tenm2. Extracellular domains are colored gray, dark blue, sky blue, cyan, and palecyan for domains 1–5, respectively; transmembrane region (TM) in brown. Domain numbers and descriptions are indicated above scheme. (B) Schematic diagram of human Lphn3. Lec and Olf domain are colored orange; Hormone Receptor (HormR), GAIN and TM domains are colored gray. Splice site of Tenm2 and Lphn3 are shown as red star and black stars, respectively. The constructs that were used in structure determination were Tenm2 -SS and Lphn3 +SS. (C) Schematic diagram of the interaction network between TEN, Lphn and FLRT at the synapse. TEN and FLRT are localized on the presynaptic cell membrane, while Lphn is localized on the postsynaptic membrane. Tenm2 -SS isoform (empty red star) forms trans-cellular complexes with Lphn3 and induces excitatory post-synaptic specializations when Lphn3 simultaneously binds to FLRT3 (left). In contrast, Tenm2 +SS isoform (filled red star) induces inhibitory post-synaptic specializations independent of Tenm2/Lphn3 interaction or FLRT3 binding (right). (D-F) The structure of the Tenm2/Lphn3 complex as obtained from single-particle cryo-EM analysis. The Lec domain of Lphn3 binds to the side of the beta-barrel of TEN. The alternatively spliced site within the b-propeller domain is distal to the Lec binding site on TEN.

The large ECRs of Tenm’s and Lphn’s form a tight trans-cellular adhesion complex (Boucard et al., 2014; Levine et al., 1994; Silva et al., 2011). The most N-terminal Lectin (Lec) and Olfactomedin (Olf) domains of Lphn interacts with Tenm2, with the Lec domain contributing most of the binding affinity (Fig. 1B). A four or five amino acid splice insert (MEQK or KVEQK) between these domains decreases the affinity of the TEN/Lphn interaction (Boucard et al., 2014). In addition to Tenm’s, Lphn’s form trans-cellular interactions also with homodimeric cell adhesion molecules called fibronectin leucine rich repeat transmembrane proteins (FLRTs), which further interact with Uncoordinated5 (UNC5s) (O’Sullivan et al., 2012) (Fig. 1C) (Sugita et al., 1998). The Lphn3/Tenm2 interaction as well as the Lphn3/FLRT3 interaction were individually reported to be important for synapse formation and organization (Boucard et al., 2014; O’Sullivan et al., 2012; Silva et al., 2011). Recent work showed that in transgenic mice in vivo, postsynaptic Lphn3 promotes excitatory synapse formation by simultaneously binding to Tenm and FLRT, two unrelated presynaptic ligands, which is required for formation of synaptic inputs at specific dendritic localizations (Sando et al., 2019) (Fig. 1C, left). Conversely, Lphn3 deletion had no effect on inhibitory synapse formation (Sando et al., 2019). The precise molecular mechanisms of neither excitatory nor inhibitory synapse formation are known.

In addition to the critical role of coincident Tenm2 and FLRT3 binding to Lphn3 for specification of excitatory synapses, alternative splicing of Tenm2 also plays a crucial role in specifying excitatory vs. inhibitory synapses (Li et al., 2018). An alternatively spliced seven-residue region (NKEFKHS) within Tenm2 acts as a switch to regulate trans-cellular adhesion of Tenm2 with Lphn’s and to induce different types of synapses in vitro (Li et al., 2018)(Fig. 1C). The Tenm2 -SS splice variant that lacks the splice insert can bind to Lphn3 in *trans* (Fig. 1C, left side). However, Tenm2 +SS, the splice variant that includes the seven amino acids is unable to interact with Lphn3 in *trans* in identical experiments (Fig. 1C, right side) (Li et al., 2018). Similarly, the same alternatively spliced site also may regulate Tenm2 trans-homodimerization (Berns et al., 2018), although no such trans-homodimerization could be detected in some assays (Boucard et al., 2014). However, the molecular mechanism of how alternative splicing regulates ligand interactions is unclear.

In agreement with in vivo transgenic mice experiments, only the splice variant of Tenm2 that can interact with Lphn3 (Tenm2 -SS) was able to promote excitatory synapses when co-expressed with FLRT3 in cultured neurons in vitro (Fig. 1C, left) (Sando et al., 2019). The other Tenm2 splice variant that cannot interact with Lphn3 (Tenm2 +SS) did not promote excitatory synapse formation when expressed alone or co-expressed with FLRT3. Instead, this Tenm2 isoform induced inhibitory post-synaptic specifications in a Lphn-independent manner likely by interacting with other unknown ligands, suggesting that alternative splicing of Tenm2 regulates excitatory vs. inhibitory synapse specification (Li et al., 2018) (Fig. 1C, right). These results indicate a multi-level coincidence signaling mechanism for the specification of synaptic connections that requires the presence of the proper combination of molecules and their appropriate alternatively spliced isoforms to colocalize in order to induce the formation of a specific type of synapse. However, the molecular details of the Tenm/Lphn/FLRT interaction are not known. Furthermore, the structural basis for the lack of the Tenm2/Lphn3 *trans* interaction in the presence of a short splice insert in Tenm2 is unclear.

Here, we have determined the 2.9 Å resolution cryo-EM structure of the Tenm2/Lphn3 complex and described the direct and simultaneous interaction of Lphn3 with both Tenm2 and FLRT3. The Tenm2/Lphn3 complex structure revealed that the N-terminal Lec domain of Lphn3 interacts with the beta-barrel domain of Tenm2. Both the Lec and the preceding Olf domains of Lphn3 face away from the alternatively spliced site within the beta-propeller domain of Tenm2, indeed providing no explanation for how the short splice insert may regulate Tenm2/Lphn3 interaction. Using a series of experimental setups that mimic either *trans*-cellular interactions between opposing cell-membranes, or *cis*-interactions in solution, we showed that alternative splicing of Tenm2 indirectly regulates the Tenm2/Lphn3 interaction by altering the accessibility of the Lphn-binding site on Tenm2 with the help of membranes, rather than directly interfering with the Lphn-binding site. Mutagenesis of the Lphn-binding site on Tenm2 abolished the Tenm2/Lphn3 interaction and had a severe and specific effect on excitatory synapse formation, but had no effect on inhibitory synapse formation. These results provide a molecular and mechanistic understanding of the multi-level coincidence binding mechanism that mediates specificity in synapse formation and circuit-wiring.

## Results

### Structure of the Tenm2/Lphn3 complex

To determine the structure of the Tenm2/Lphn3 complex, the ECR of human Tenm2 lacking the EGF repeats that are responsible for *cis*-dimerization (Tenm2 -SS ECRΔ1, encoding residues 727-2648, Fig. 1A) and the full ECR of human Lphn3 (Lphn3 +SS ECR, encoding residues S21-V866, Fig. 1B) were co-expressed in insect cells using the baculovirus expression system. The complex was purified by size-exclusion chromatography and the complex structure was determined by single particle cryo-EM. After multiple rounds of 3D classification, two cryo-EM maps were obtained: One corresponding to the monomeric Tenm2 -SS ECRΔ1 in complex with Lphn3 ECR at a nominal resolution of 2.9 Å (from 9.7% of particles, Figure S2 and S4C), and the other corresponding to the monomeric Tenm2 ECRΔ1 with a better resolved β-propeller domain at a nominal resolution of 3.0 Å (from 3.8% of particles, Figure S3 and S4D). A near-atomic resolution model of the protein complex was built using the available Tenm2 structure (PDB: 6CMX) and the Lec Olf structure (PDB:5AFB) (Figure 1D, Figure S1-4, Table 1).

**Table 1.** Cryo-EM data collection, refinement and validation statistics.

|  | #1 TEN2/LPHN3 complex<br>(EMD-21205)<br>(PDB 6VHH) | #2 TEN2<br>(EMD-21205) |
| --- | --- | --- |
| **Data collection and processing** |  |  |
| Magnification | 81,000 | 81,000 |
| Voltage (kV) | 300 | 300 |
| Electron exposure (e-/Å<sup>2</sup>) | 60.1 | 60.1 |
| Defocus range (µm) | -1.0 to -2.5 | -1.0 to -2.5 |
| Pixel size (Å) | 0.54 | 0.54 |
| Symmetry imposed | C1 | C1 |
| Initial particle images (no.) | 4,475,958 | 4,475,958 |
| Final particle images (no.) | 436,208 | 170,133 |
| Map resolution (Å) | 2.97 | 3.07 |
| FSC threshold | 0.143 | 0.143 |
| Map resolution range (Å) | 2.4-4.5 | 2.4-4.5 |
| **Refinement** |  |  |
| Initial model used (PDB code) | 6CMX |  |
| Model resolution (Å) | 3.0 |  |
| FSC threshold | 0.5 |  |
| Model resolution range (Å) |  |  |
| Map sharpening *B* factor (Å<sup>2</sup>) |  |  |
| Model composition |  |  |
| Non-hydrogen atoms | 14271 |  |
| Protein residues | 1821 |  |
| Ligands | BMA:5<br>NAG: 14 |  |
| *B* factors (Å<sup>2</sup>) |  |  |
| Protein | 53 |  |
| Ligand | 67 |  |
| R.m.s. deviations |  |  |
| Bond lengths (Å) | 0.015 |  |
| Bond angles (°) | 1.034 |  |
| Validation |  |  |
| MolProbity score | 2.02 |  |
| Clashscore | 10.16 |  |
| Poor rotamers (%) | 0.61 |  |
| Ramachandran plot |  |  |
| Favored (%) | 91.78 |  |
| Allowed (%) | 8.05 |  |
| Disallowed (%) | 0.17 |  |

Our Tenm2/Lphn3 complex structure comprises a heterodimer of ∼100 × 50 × 115 Å. The Lec domain of Lphn3 interacts with the side of the barrel domain of Tenm2 (Figure 1D-F). The Tenm2 ECR is assembled as a large cylindrical barrel sealed by the β-propeller and Ig-like domains at the bottom; and the toxin-like domain protrudes from and attaches to the side of the barrel as previously reported (Jackson et al., 2018; Li et al., 2018). The Lec domain of Lphn3 and the toxin-like domain of Tenm2 bind to opposite faces of the beta-barrel in a seemingly parallel orientation to each other (Figure 1D-F). Apart from sidechain rotamer changes, no major conformational changes are observed when the complex structure is compared with the individual structures of Tenm2 or Lphn3. Nine N-linked glycosylation sites (on residues N1490, N1586, N1647, N1681, N1766, N1867, N2071, N2211, N2522.) and five disulfide bonds (C1394-C1402), (C1396-C1404), (C1106-C1109), (C1210-C1218), (C1277-C1330) were observed in Tenm2, while four disulfide bonds were detected in Lphn3 (C36-C66), (C78-C110), (C91-C97), (C45-C123).

Analysis of the cryo-EM maps at a lower threshold also revealed continuous a density for the Olf domain that extended from the Lec domain towards the opposite side from the β-propeller domain of Tenm2 (Figure 2A). In spite of the lower resolution, it was possible to fit the available Lphn3 Olf domain structure in this density (Figure 2B). The Olf domain is positioned in close proximity to the top of the Tenm2 beta-barrel, although it is not in contact with Tenm2 (Figure 2B). The presence of the splice insert (KVEQK) between the Lec and OLF domains in our Lphn3 construct likely causes the lack of interaction of OLF domain with TEN. The remaining C-terminal domains of Lphn3 (the hormone receptor and GAIN domains) for which there is no EM density likely extend from the opposite side, away from the Tenm2 TM domain located at the Tenm2 N-terminus. This orientation positions the membrane-anchored TM domain of Lphn3 on the opposite side from the membrane-anchored TM domain of Tenm, and thus is compatible with a *trans*-cellular interaction of Tenm with Lphn (Figure 2C).

**Figure 2:**
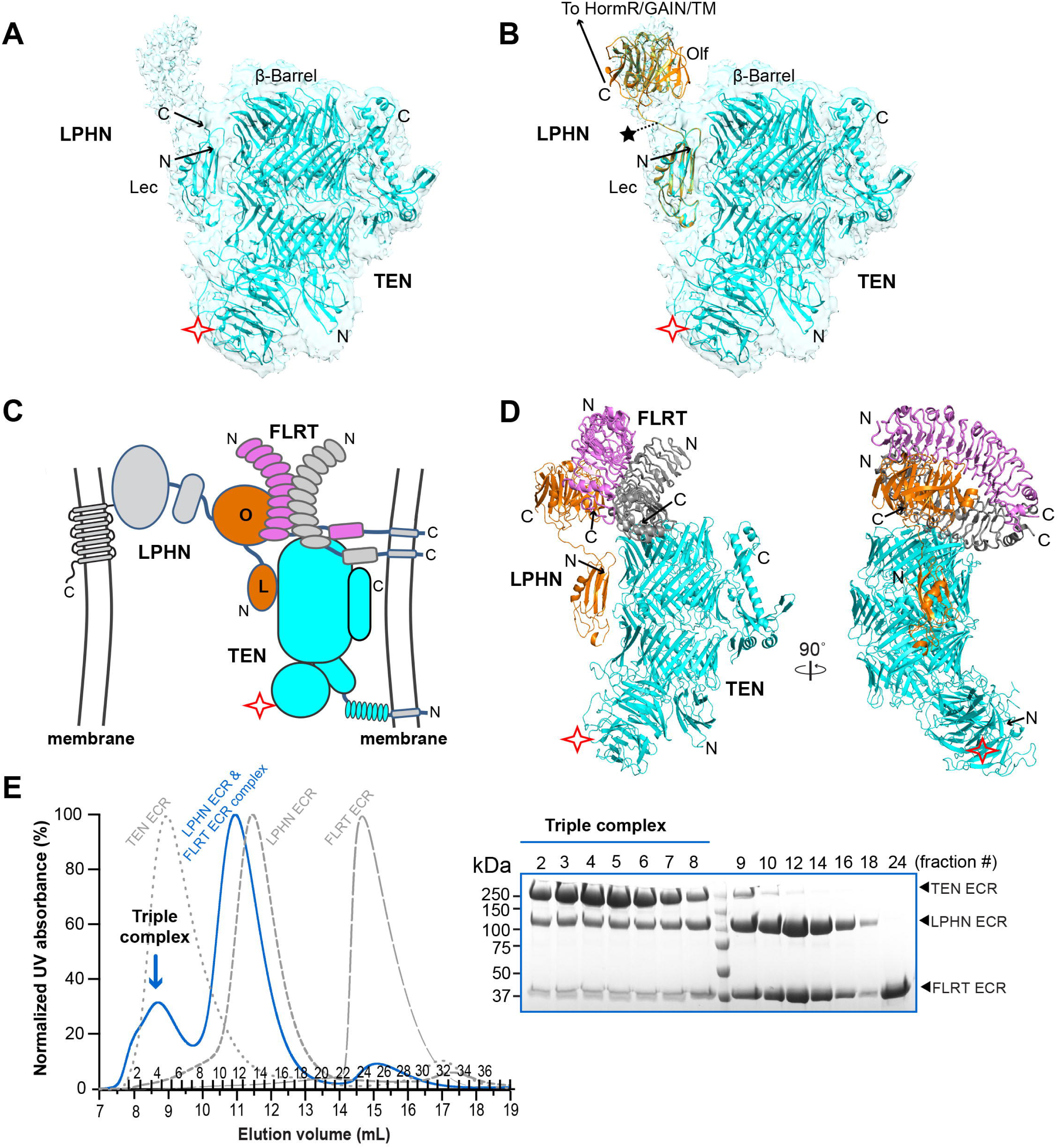
Lphn3 interacts with Tenm2 and FLRT3 simultaneously. (A) Continuous density C-terminal to the Lec domain of Lphn3 revealed by analysis of the cryo-EM maps at a lower threshold. (B) Manual fitting of the Olf domain from the Lphn3 Lec-Olf structure (PDB: 5AFB) into the extra EM density. Ribbon diagram of TEN3/Lphn3-Lec complex is colored cyan and the Lphn3 Lec-Olf domains are colored orange. Lec domains are superimposed. (C) Schematic diagram of the trans-cellular positioning of the Tenm2/Lphn3 complex; and the formation of a ternary complex between Tenm2, Lphn3 and FLRT3. (D) Superimposition of the TEN3/Lphn3-Lec-Olf complex with the Lphn3/FLRT3 complex structure (PDB: 5CMN). The Olf domains are superimposed. Tenm2 and Lphn3 are colored cyan and orange, respectively. FLRT3 molecules in the FLRT3 dimer are colored magenta and gray. The second Lphn3 molecule that might be bound to the FLRT3 dimer is not shown. An UNC5 molecule that might be bound to the magenta FLRT3 is not shown. (E) SD200 size-exclusion chromatography profile of the Tenm2/Lphn3/FLRT3 complex (Blue line) as compared to the profiles of individual proteins (dashed gray lines) showing that Lphn3 binds to Tenm2 and FLRT3, simultaneously. Size-exclusion fractions are run on an SDS-PAGE gel. Triple complex is indicated by blue arrow.

### Lphn3 binds to both Tenm2 and FLRT3 simultaneously and forms a trimeric complex

Lphn3 proteins are involved in heterodimeric interactions with Tenm’s and FLRTs and coincidence binding of both FLRTs and Tenm’s is required for excitatory synapse formation (Figure 1C, left). Additionally, FLRTs interact with UNC5s and form homodimers that are incompatible with their UNC5 binding. However, whether these interactions are compatible is unclear. Thus, we investigated whether the Tenm2, Lphn3 and FLRT3 interactions are compatible with each other, or in other words, whether Tenm2, Lphn3, and FLRT3 can form a trimeric complex. The availability of the Lphn3/FLRT3 complex structures and of the Lphn3 Lec-Olf structure enabled us to compare structures, and to predict and test the compatibility of the possible interactions of Tenm, Lphn3 and FLRT3 with each other. Intriguingly, superimposition of the Lec domain from the Lphn3/FLRT3 structure with the Lec domain from the Lphn3/Tenm2 complex structure showed that FLRT3 and Tenm2 bind to distinct domains on Lphn3 and that there are no clashes between Tenm2 and FLRT3, suggesting that Lphn3 can simultaneously bind to Tenm2 and FLRT3 (Figure 2C, D).

In order to test whether this model is correct, we co-expressed recombinant FLRT3 LRR, Lphn3 full ECR, and Tenm2 -SS full ECR in insect cells, purified the complex by gel filtration chromatography and analyzed the fractions by SDS-PAGE (Figure 2E). Both Lphn3 ECR and FLRT3 LRR elution volumes shifted to the left as compared to the elution volumes of the isolated FLRT3 and the isolated Lphn3. All three proteins eluted in the same fractions, indicating the formation of a trimeric Tenm2/Lphn3/FLRT3 complex (Figure 2E). These results suggest that Tenm2 and FLRT3, both ligands of Lphn3, can simultaneously bind to Lphn3 and form a trimeric complex in vitro, supporting the in vivo observations that coincident binding of both Tenm2 and FLRT3 to Lphn3 is required for excitatory synapse formation (Sando et al., 2019).

### The binding interface of the Tenm2/Lphn3 complex is conserved

To visualize conserved and variable regions of the Tenm2 β-barrel domain and the Lphn3 Lec/Olf domains, we mapped the conservation of residues on the Tenm2/Lphn3 complex structure, and colored residues from most conserved (magenta) to least conserved (cyan) (Fig. 3A). The interaction surfaces of Tenm’s and Lphn’s correspond to one of the most conserved regions (yellow ovals in Fig. 3A). Specifically, residues on the Tenm2 β-barrel, on the Lec domain and also on the OLF domain that face the binding interface are highly conserved. As the Tenm2 β-barrel is homologous to bacterial Tc-toxins, we also analyzed the conservation between bacterial toxins by mapping the conservation of residues between bacterial toxins on the homologous bacterial TcC toxin structure (PDB ID: 4O9X) (Meusch et al., 2014) and observed that the identical surface of bacterial Tc toxins is not conserved (Fig. S5A). Our cryo-EM map thus revealed that Lphn3 binds to a highly conserved surface on the barrel of Tenm2 that likely evolved to bind to Lphn’s after diverging from bacterial toxins.

**Figure 3.**
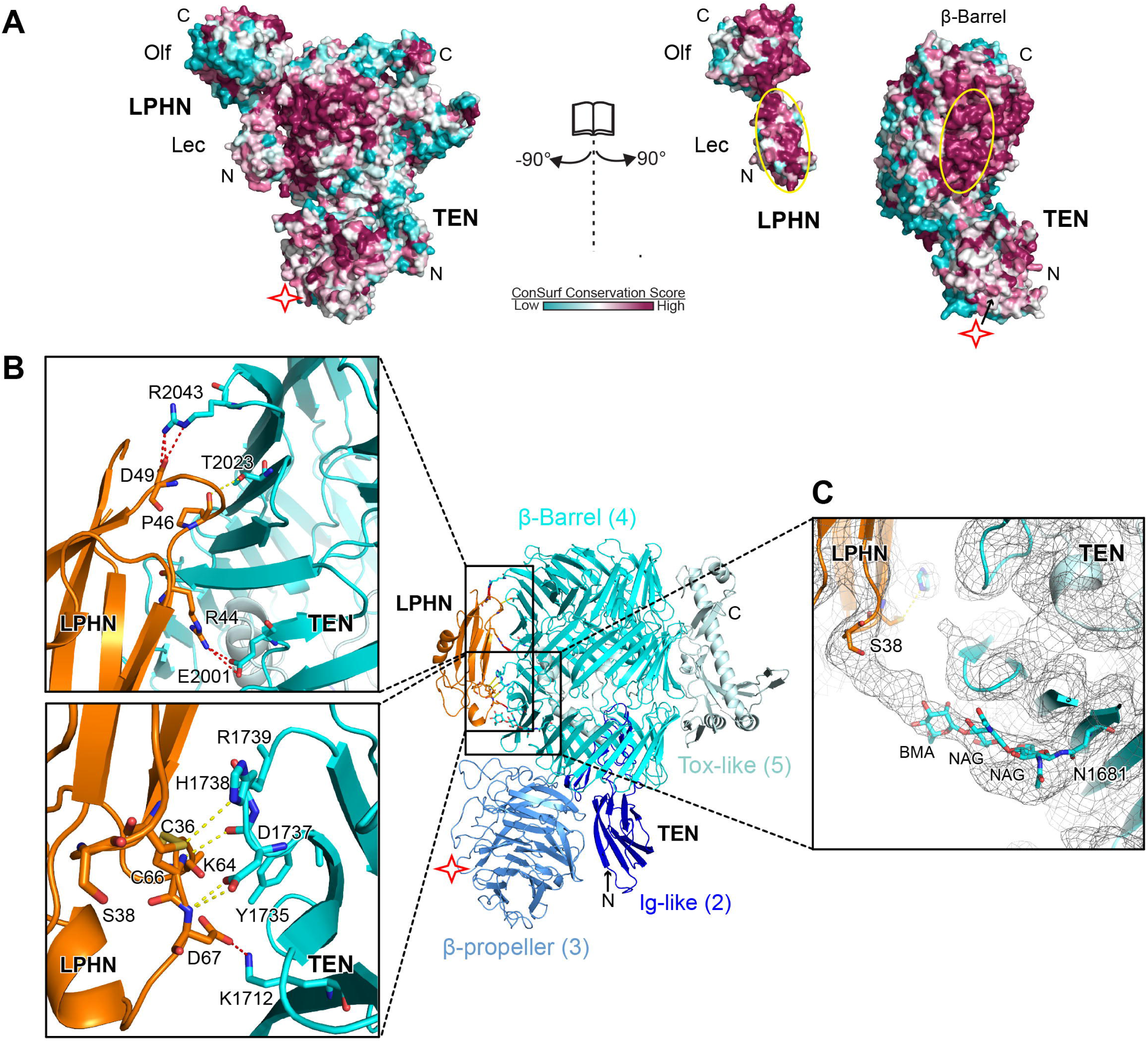
Tenm2 and Lphn3 interaction is mediated by conserved residues. (A) The TEN/Lphn3 binding interface is conserved. The structure of the Tenm2/Lphn3 complex is shown in surface representation on which the conservation of residues is mapped from most conserved (magenta) to least conserved (cyan) (using the ConSurf server; Landau et al., 2005). The Lphn-binding site on Tenm2 and the Tenm2 -binding site on Lphn3 are indicated by yellow circles. (B) Ribbon diagram of the Tenm2/Lphn3 heterodimer showing the interface between Tenm2 and Lphn. Close-up view of the binding interface shows residues involved hydrogen bonds and salt bridge shown by yellow dashes and red dashes, respectively. The Tenm2 mutation “DHR” mutation (D1737N, H1738T, R1739T) which disrupt the Tenm2/Lphn3 are shown as sticks. (C) Close-up view for the EM density showing the close packing of the conserved N-linked glycosylation on Tenm2-N1681 with Lphn3-S38 on the Lec domain.

The Lec domain of Lphn3 belongs to the sea urchin egg lectin (SUEL) related Lec family. It adopts a kidney shape with dimensions of 20Å × 20Å × 50Å, and is composed of five beta strands and a single alpha helix, interconnected by four conserved disulfide bonds (Vakonakis et al., 2008). The docking of the complementary surfaces of the beta-sandwich of the Lphn3 Lec domain to the concave surface of the Tenm2 beta barrel creates an average interface area of 690 Å^2^. The high affinity of the Tenm2/ Lphn3 complex is achieved by a combination of interactions, comprised of salt bridges, hydrogen bonds, and long-range electrostatic interactions (Figure 3B). Notably, salt bridges located at the top, middle and the bottom of the interface stabilize the interaction (two salt bridges between ARG44 NH1 and GLU2001 OE1/OE2; two between ASP49 OD1 and ARG2043 NH2/NE; one between ASP49 OD2 and ARG2043 NH2; and one between ASP67 OD1 and LYS1712 NZ)\ (Fig. 3B). The extensive network of salt bridges likely helps achieve the high affinity of the Tenm2 /Lphn3 complex. In addition, two residues on Tenm2 (D1737 and H1738) play important roles by interacting with a disulfide bond (C36, C66) and D67 of the Lec domain (Fig. 3B).

Interestingly, the cryo-EM map showed clear density of the Lec domain interacting with a glycochain originating from glycosylation at N1681 (Fig. 3C). The glycochain inserts into a well conserved sugar-binding pocket of the SUEL-related Lec domain of Lphn3 (Fig. S5B), suggesting that in contrast to previous notions (Vakonakis et al., 2008), the Lec domain of Lphn3 is still capable of binding carbohydrates. Due to the uncertainty of the glycochain topology, we only modeled the first three sugars of N-glycosylation (NAG-NAG-BMA) into the density. Nevertheless, it is clear that the Lec domain is interacting with the glycochain of Tenm.

In order to specifically abolish the interaction of Tenm2 with Lphn3 without interfering with other interactions or the cell-surface localization of Tenm2, and to confirm the validity of the binding interface that we observed in the Tenm2/Lphn3 complex structure, we designed surface mutations on full-length Tenm2 that change only a few atoms on the protein surface. Several Tenm2 mutations were designed, including the “DHR” (D1737N, H1738T, R1739T) mutation that alters residues at the Lphn3 Lec domain binding interface (Fig. 3B). To ensure that the mutant proteins are properly folded, we first examined the expression levels and surface transport of all Tenm2 mutants. We eliminated misfolded mutants that are likely to be poorly expressed and unable to reach the cell surface and exclusively used mutants that had no localization problems (Fig. S5C). All constructs included extracellular C-terminal Flag and HA tags to allow for measurements of expression levels and of the cell-surface localization of WT and mutant proteins. HEK293T cells transfected with WT and mutant Tenm2 constructs were stained without detergent permeabilization (to label only the cell-surface localized protein) with an antibody suitable to react with the extracellular tag on the proteins, and the amount of surface-exposed Tenm2 was assessed by indirect immunofluorescence using flow cytometry and immunostaining. The properly trafficked mutants were tested for their ability to interact with Lphn3 (see below).

### Membranes enforce a splice variant-dependent docking geometry on the *trans*-cellular Tenm2/Lphn3 complex

A seven-amino acid alternatively spliced site on the beta-propeller domain of Tenm2 regulates the Tenm2/Lphn3 interaction, and, consequently, excitatory vs. inhibitory synapse formation (Li et al., 2018). A striking observation from the Tenm2/Lphn3 complex structure was that the Lphn3 binding site on Tenm2 is located distal to the alternatively spliced sequence in the Tenm2 β-propeller. This observation is very surprising because in other ligand interactions regulated by alternative splicing, the alternatively spliced sequence is usually located at the ligand-binding interface (Arac et al., 2007; Wilson et al., 2019). Thus, the Tenm2/Lphn3 interaction is the first case to our best knowledge in which alternative splicing regulates a ligand interaction remotely.

We hypothesized that the membranes of two opposing cells may impose a docking geometry on Tenm2 and Lphn3 that is critical for their *trans*-cellular adhesion in the extracellular space between the membranes, and that alternative splicing might control the docking geometry of Tenm2. Consequently, with an altered geometry, Lphn3 may not be able to access its binding site on Tenm. This hypothesis suggests that insertion or deletion of the alternatively spliced sequence does not affect the Lphn3 binding surface on Tenm. It also suggests that when one or more membranes are removed from the experimental system, Lphn3 and Tenm would interact with each other independent of alternative splicing because they will not be restricted by the membranes to which they are anchored. In order to test this hypothesis, we used various experimental setups in which the restraints that act on the docking geometries of Tenm2 and Lphn3 vary from high to low (Figure 4A, from 1 to 3).

**Figure 4.**
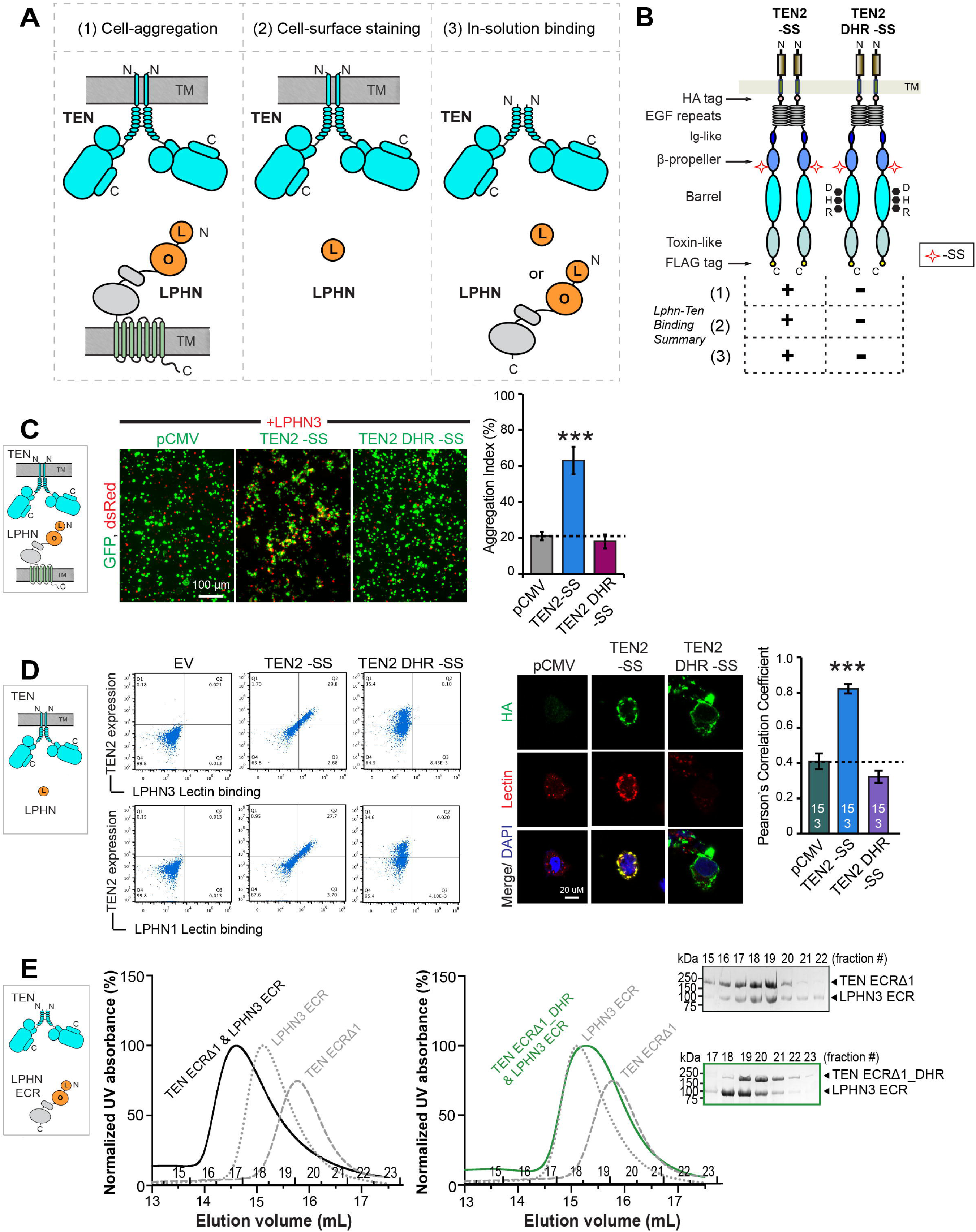
Binding site mutations on Tenm2 abolish Lphn3 binding in both trans and cis. (A) Three experimental setups (1-3) with decreasing restraints on the docking geometry of Lphn3 and Tenm2 during their interaction. (A1) Setup for trans-cellular interaction of full-length Tenm2 with full-length Lphn3 in cell-aggregation experiments. Both proteins are anchored on the cell-membranes and their mobility is restricted by their lateral diffusion within the membrane; (A2) Setup for cis-interaction of full-length Tenm2 and soluble biotinylated Lec domain of Lphn3 in either flow cytometry or cell surface staining experiments. Tenm2 is anchored on the cell-membrane, but the Lec domain of Lphn3 freely rotates in solution; (A3) Setup for in-solution experiments for cis-interaction of soluble Tenm2 and soluble Lphn3 in the absence of any membranes. (B) Diagram for WT Tenm2 -SS and Tenm2 DHR -SS constructs that were used in the below experiments. “DHR” mutation (D1737N, H1738T, R1739T) is on the Tenm2 β barrel located at the Lphn3 binding interface (red stars). Results for the interaction of Tenm2 and Lphn3 in different experimental setups (A1-3) are summarized in the table. The DHR mutation breaks the interaction of Tenm2 with Lphn3 in all experimental setups. (C) Representative images for cell-aggregation assays with WT Tenm2 -SS or Tenm2 DHR -SS and full-length Lphn3. WT Tenm2 -SS induces cell aggregation with Lphn3, while Tenm2 DHR -SS abolishes cell aggregation. HEK293 cells were co-transfected with the indicated Tenm2 or Lphn3 and either tdTomato or EGFP as indicated. Scale bar indicates 100 µm. All experiments were performed in three independent culture preparations. Quantification of aggregation index (%) in indicated conditions is shown on the right (n = 3 independent replicates with quantifications from 3 randomly selected imaging fields per replicate; ***p < 0.001 by ANOVA). (D) WT Tenm2 -SS and Tenm2 DHR -SS expressed in mammalian cells were tested for their ability to bind soluble biotinylated Lphn3 or Lphn1 Lec domain using flow cytometry experiments (left) and using cell surface staining assays (right). The “DHR” mutation abolish the cis interaction between Tenm2 and Lphn. Tenm2 construct expression was determined by HA tag fluorescence (Y axis) and purified Lec binding to Tenm2-expressing cells was measured by fluorescence of DyLight attached to neutravidin (X axis). Dot plots represent the correlation between Tenm2 expression and Lphn3 binding in Tenm2-transfected cells and empty vector-transfected control. Black cross on the plot shows the ‘‘high Tenm2 expression and high Lphn3 binding’’ gate. Scale bar indicates 20 µm. Quantification of cell surface binding assays are shown next to the image. (E) Size-exclusion chromatograms showing the formation of a binary complex between soluble Tenm2 -SS ECRd1 and full Lphn3 ECR (left, black line). Elution profile for individual Tenm2 -SS ECRd1 and individual Lphn3 ECR are shown for reference (dotted lines). Tenm2 -SS ECRd1 DHR mutant does not bind Lphn3 ECR (right, green line), as observed by the lack of co-elution in fractions ran on SDS-PAGE gel. Colors of the chromatograms match the colors of box around the SDS-PAGE gel.

First, we conducted cell-aggregation assays with HEK293 cells in which a population of HEK cells expressing full-length Tenm2 are mixed with a different population of HEK cells expressing full-length Lphn3, and cell aggregation is monitored as a function of Tenm2/Lphn3 interaction (Fig. 4A(1)). These experiments mimic *trans*-cellular interaction as they detect the binding of two full-length proteins that are anchored on opposing cell membranes (we refer to these experiments as “*trans*” hereon). Cell-aggregation experiments apply high restraints on the docking geometries of the proteins because both proteins can diffuse only laterally in two dimensions within the plane of the membrane bilayer. The cell-aggregation experiments are the best imitation for the in vivo interaction of Tenm2 and Lphn3, where full-length proteins are on the cell surfaces of neighboring cells during development or synapse formation. Second, we used flow cytometry experiments and cell-surface staining experiments in which the binding of a soluble protein to a membrane-anchored protein is tested (we refer to these experiments “*cis*” hereon, although it should not be confused with other commonly used meanings of *cis*) (Fig. 4A(2)). Soluble fragments of the Lphn3 ECR were tested for their ability to bind to HEK293T cells expressing full-length Tenm2 on the cell surface. These experiments apply intermediate restraints on the docking geometries of the proteins because the membrane-anchored Tenm2 can diffuse only laterally in two dimensions within the plane of membrane bilayer, while the soluble Lphn3 ECR fragment can freely diffuse in three dimensions in solution (Fig. 4A(2)). Third, we used size exclusion chromatography in which the binding of two soluble proteins experience no restraints. Here, binding of the ECRs of Tenm2 and Lphn3 is tested in the absence of any membranes (we use the term “*cis*” for these experiments as well) (Fig. 4A(3)). Importantly, any intrinsic restraints that may be originating from the intrinsic conformation of the proteins may still act on any of the above experiments. We used a combination of these experimental setups to investigate the effect of various restraints on the ability of Tenm2 to interact with Lphn3.

We examined two sets of Tenm2 constructs in these experimental setups: (i) Tenm2 –SS carrying Lphn3 binding site mutations (Tenm2 –SS DHR) was compared to WT Tenm2 -SS to observe the effect of Lphn-binding site mutations on TEN/Lphn3 interaction (Fig. 4B). We expect that this mutant should abolish TEN/Lphn3 interaction in all experimental setups because it directly disrupts the Lphn-binding site on TEN. (ii) WT Tenm2 +SS was compared to WT Tenm2 –SS to observe the effect of inclusion of the splice insert on TEN/Lphn3 interaction (Fig. 5A). We expect that, if the inclusion of the alternative splice insert is disrupting the binding interface on Tenm2 for Lphn, then Tenm2 +SS variant should not bind Lphn3 in any of the experimental setups. However, if the insertion of the alternative splice insert is acting by a different mechanism, such as changing the docking geometry of Tenm2 onto Lphn, then, Tenm2 +SS isoform may bind Lphn3 in “*cis*” setups where binding restraints on Tenm2 and Lphn3 are relaxed (Fig. 4A(2 and 3)).

**Figure 5.**
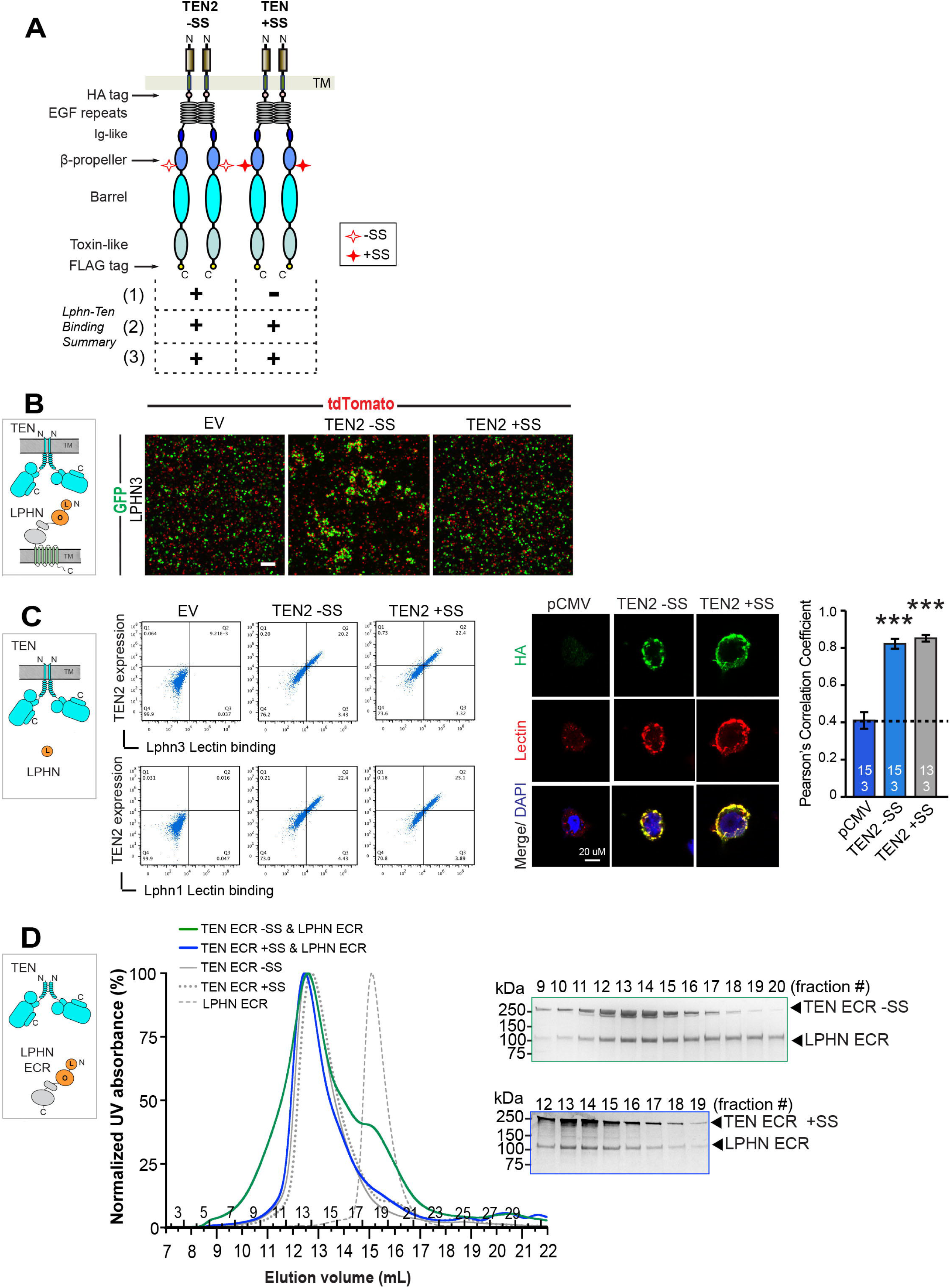
Membrane anchoring restricts alternative splice-dependent interaction of Tenm2 to Lphn3. Same three experimental setups as in Figure 4A(1-3) were used to test the effect of alternative splicing on Tenm2/Lphn3 interaction. Figure outline is identical in principle to that of in Figure 4. (A) Diagram for WT Tenm2 -SS and WT Tenm2 +SS constructs that were used in the below experiments. The seven-amino acid splice site on the Tenm2 β-propeller is indicated by empty or filled red stars. Results for the interaction of Tenm2 and Lphn3 in different experimental setups (as in Figure 4A1-3 and Figure 5B, C, D) are summarized in the table. The insertion of the splice site breaks the interaction of Tenm2 with Lphn3 only in the cell-aggregation assays, but not in the other experimental setups. (B) Representative images for cell-aggregation assays with Tenm2 -SS or Tenm2 +SS and full-length Lphn3. Tenm2 -SS induces cell aggregation with Lphn3, while Tenm2 +SS abolishes cell aggregation. (C) Tenm2 -SS and Tenm2 +SS expressed in mammalian cells were tested for their ability to bind soluble biotinylated Lphn3 or Lphn1 Lec domain using flow cytometry experiments (left) and using cell surface staining assays (right). Both -SS and +SS mediate the interaction between Tenm2 and Lphn in *cis*. Quantification of cell surface binding assays are shown next to the image. The cell-surface staining assays in Fig. 5C was performed in the same experiment as in Fig. 4D, and thus the control images are identical. (D) Size-exclusion chromatograms showing the formation of binary complexes between soluble full Tenm2 ECR and full Lphn3 ECR (left, blue and green lines). Elution profile for individual Tenm2 -SS ECR, Tenm2 +SS ECR and Lphn3 ECR are shown for reference (grey lines). Both Tenm2 -SS ECR and Tenm2 +SS ECR bind to Lphn3 ECR (green and blue lines, respectively), as also observed by co-elution in the fractions ran on SDS-PAGE gel. Colors of the chromatograms match the colors of box around the SDS-PAGE gel.

The first set of experiments testing the effect of point mutations on the Lphn3 binding surface showed that Tenm2 -SS DHR mutant was unable to bind to Lphn3 in all experimental setups, including cell-aggregation experiments (Fig. 4C), flow cytometry experiments (Fig. 4D, left), cell-surface staining experiments (Fig. 4D, right) and gel filtration experiments (Fig. 4E). These results show that this mutation destroys the binding interface on Tenm2 for Lphn3 and abolishes complex formation in *trans* and *cis* (Fig. 4). The second set of experiments testing the effect of the alternatively spliced sequence of Tenm2 on Lphn3 binding, however, displayed differential effects in *trans* and *cis* experimental setups (Fig. 5). In cell-aggregation experiments, full-length Tenm2 lacking the β-propeller splice insert robustly induced *trans*-cellular aggregation with full-length Lphn3 (Fig. 5B). Intriguingly, as we previously showed, inclusion of the seven-amino-acid splice insert in the β-propeller eliminated trans-cellular adhesion with Lphn3 (Fig. 5B). In flow cytometry experiments, however, the affinity of Tenm2 for the soluble LEC domain (that is unattached to the membranes) was not affected by inserting the seven-residue segment (Fig. 5C, left). Cell-surface staining experiments also showed that the soluble LEC domain of Lphn3 binds to both Tenm2 splice isoforms, confirming the flow cytometry experiments (Fig. 5C, right). Finally, gel filtration chromatography experiments performed with the full-length Tenm2 and Lphn3 ECRs as soluble proteins unattached to membranes showed that when no restraints are applied on Tenm2 and Lphn3, both Tenm2 splice isoforms robustly interacted with Lphn3 (Fig. 5D). Thus, we suggest that *trans*-cellular Tenm2-Lphn3 interactions are regulated by β-propeller alternative splicing likely due to conformational restrains in the context of full-length proteins.

### Mutations at the Tenm2/Lphn3 binding interface affect splice variant-dependent excitatory but not inhibitory synapse specification

Previous work showed that the splice isoforms of Tenm2 (Tenm2 +SS and Tenm2 -SS) induce different synaptic specifications in artificial synapse formation assays. In these assays, HEK293 cells expressing Tenm2 variants were co-cultured with primary neurons; and inhibitory and excitatory synapse formation was monitored for pre- and post-synaptic differentiation for both types of synapses (Li et al., 2018). The results showed that Tenm2 +SS induced GABAergic (inhibitory) postsynaptic specializations but failed to induce glutamatergic (excitatory) postsynaptic specifications (Li et al., 2018). On the other hand, initially, Tenm2 -SS failed to recruit both excitatory and inhibitory synaptic markers (Li et al., 2018). However, when FLRT3, another Lphn3 ligand that on its own is also unable to induce pre- or postsynaptic specializations, was co-expressed in HEK293 cells with the Tenm2 -SS, these molecules together potently induced excitatory but not inhibitory postsynaptic specializations (Sando et al., 2019) (Fig. 1C, left side).

As only the Tenm2 -SS isoform is capable of interacting with Lphn3 in a *trans*-configuration, we speculated that the DHR mutation that abolishes the interaction of Tenm2 with Lphn3 by demolishing the binding site should affect excitatory synapse formation, but should not impair inhibitory synapse formation that is mediated by the Tenm2 +SS isoform because inhibitory synapse formation is independent of the Tenm2/Lphn3 interaction. To test this hypothesis, we engineered the DHR mutation on both the full-length Tenm2 -SS and the Tenm2 +SS isoforms, and tested its effect on the induction of either excitatory (-SS variant) or inhibitory (+SS variant) post-synaptic specializations in the artificial synapse formation assay (Fig. 6A). As previously shown, WT Tenm2 -SS induced excitatory post-synaptic specializations when co-expressed with FLRT3 (Fig. 6B-C); and WT Tenm2 +SS induced inhibitory post-synaptic specializations (Fig. 6D-E) (Li et al., 2018; Sando et al., 2019). We observed that the DHR mutant attenuated the formation of excitatory synapses when compared with wild type Tenm2 -SS (Fig. 6B-C). However, the same mutant triggered inhibitory post-synaptic specializations similar to that of the wild type Tenm2 +SS (Fig. 6D-E), as predicted. These results indicate that Lphn3 binding mediates the excitatory synapse formation of Tenm2 -SS, whereas binding of Lphn3 to Tenm2 is not involved in inhibitory synapse formation.

**Figure 6.**
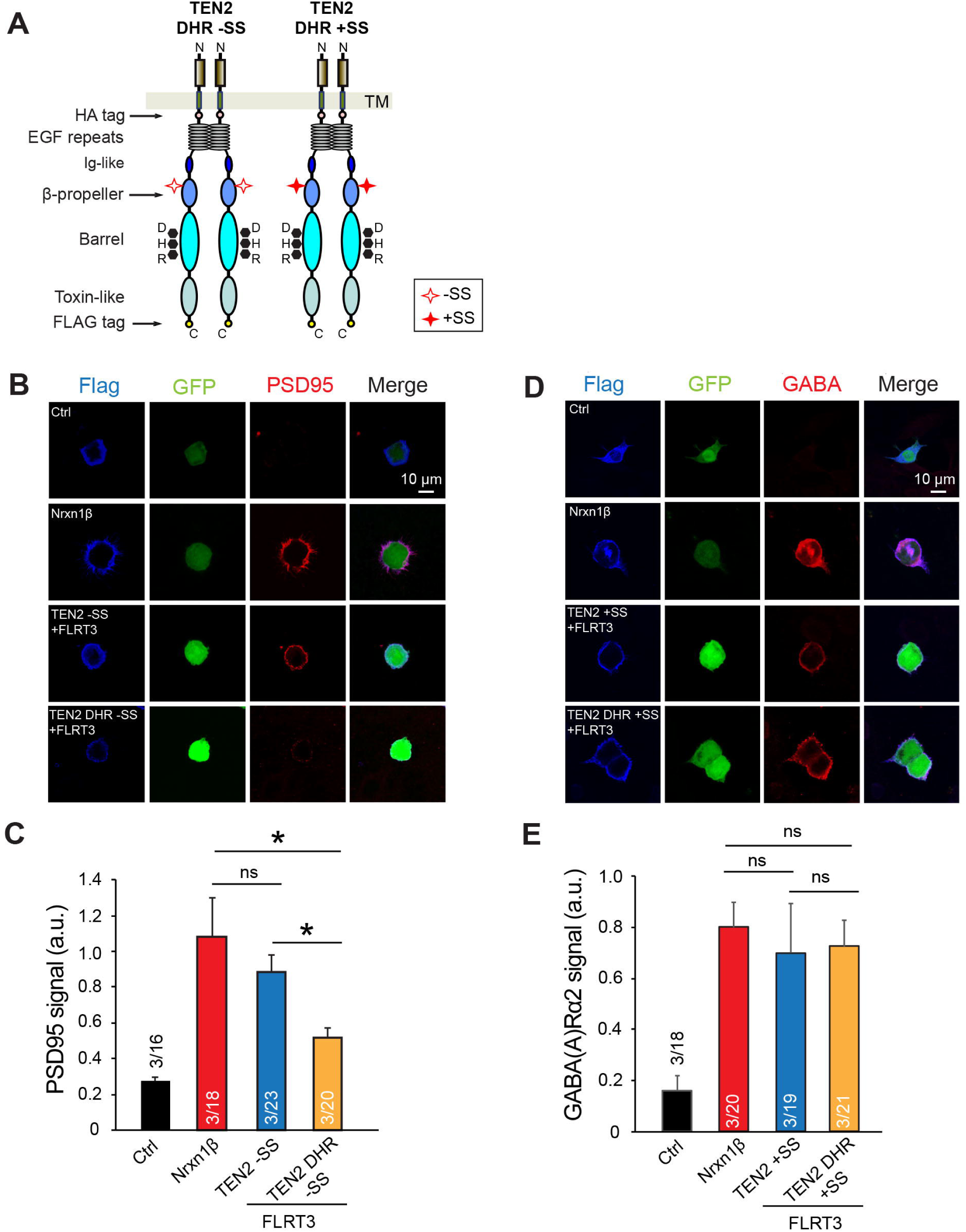
Binding site mutations on Tenm2 selectively abolish excitatory but not inhibitory synapse formation. (A) Diagram for TEN2 DHR -SS and TEN2 DHR +SS constructs that were used in the below experiments. The seven-amino acid splice site on the TEN2 β-propeller is indicated by empty or filled red stars; DHR mutation is indicated by black dots. (B-C) Artificial synapse formation assay showing that Lphn-binding mutant (DHR) of Tenm2 -SS attenuated excitatory synapse formation. HEK293T cells are co-transfected with indicated cell adhesion molecules and GFP, and co-cultured with cortical neurons. Cultures were subsequently immunostained for the excitatory postsynaptic synapse marker PSD95. Representative images (B) and quantifications of PSD95 signals (C) are shown. (D-E) Lphn-binding mutant (DHR) of Tenm2 +SS did not affect inhibitory synapse formation. Similar in B and C, except that immunostaining for the inhibitory postsynaptic synapse marker GABA(A)γ2 was performed. Data in (C) and (E) are means ± SEM from three independent experimental replicates. *P < 0.05 (one-way ANOVA).

## Discussion

Teneurins and latrophilins are multifunctional transmembrane proteins that perform important biological roles via trans-cellular interactions. The function of Lphn3 in excitatory synapse formation requires simultaneous binding of Lphn3 to both Tenm’s and FLRT’s, suggesting that a coincidence signaling mechanism mediates specificity of synaptic connections. Furthermore, synapse specificity is regulated by alternative splicing of Tenm2 because only the Lphn3-binding splice variant of Tenm2 can induce excitatory synapse formation. The variant of Tenm2 that is unable to bind Lphn3 induces inhibitory synapses in a manner that is likely Lphn3-independent. A molecular understanding of the Tenm2-Lphn3 complex and its critical regulation by alternative splicing to specify excitatory vs. inhibitory synapse specification is essential for progress in understanding synapse formation.

In this study, we determined the cryo-EM structure of the Lphn3-Tenm2 complex which revealed that the N-terminal Lec-like domain of Lphn3 binds to the side of the Tenm2 barrel opposite to the toxin-like domain (Figure 1D-F, also see Figure S6 and supplementary discussion in Figure S6 legend for comparison with previous reports). The nearby Olf-like domain of Lphn3 faces away from the N-terminus of Tenm2, positioning the membrane anchored domains of Lphn3 and Tenm2 opposite from each other, consistent with a trans-cellular, rather than cis interaction (Figure 7A). A FLRT3 molecule can simultaneously bind to the Olf-like domain of Lphn3 and form a trimeric Tenm2-Lphn3-FLRT3 complex (Figure 2C-E). It is likely that the trimer may accommodate binding of a second FLRT3 molecule to enable FLRT3 dimerization or binding of an UNC5 molecule on the FLRT monomer. However, the alternatively spliced sequence of Lphn3 in the loop between the Lec and Olf-like domains may affect the movement of the Olf-like domain and may cause clashes between the additional FLRT3 or UNC5 with Tenm2. In this scenario, FLRT3 dimerization or the FLRT3/UNC5 interaction may not be possible, leading to further rearrangement of the protein-protein interaction network at the synapse.

**Figure 7:**
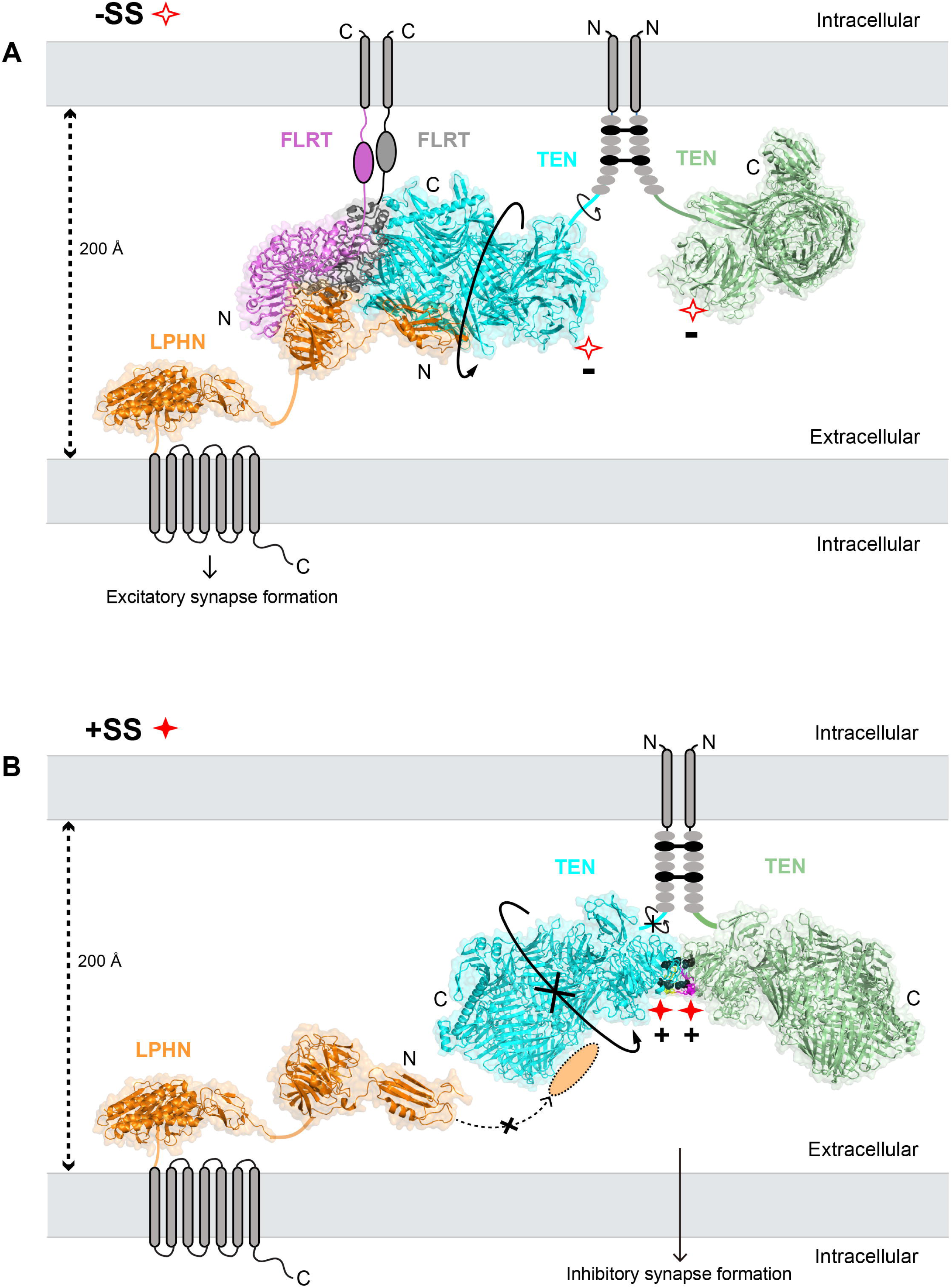
Model for the splice variant-dependent interaction of Tenm2 with Lphn3. The model depicts how alternative splicing acts as a molecular switch to determine which adhesion partner Tenm2 binds to and, accordingly, which type of synapse Tenm2 specifies. Both Tenm2 isoforms form a cis-dimer on the presynaptic membrane through two disulfide bonds formed between the 2^nd^ and 5^th^ EGF repeats (black sticks). (A) Tenm2 -SS isoform has rotational flexibility (arrows) that enables Tenm2 to find the correct docking geometry in order to bind to the Lec domain of Lphn3 expressed on the neighbor cell (Li et al., 2018). Such rotational flexibility also allows FLRT3 to bind to the Olf domain of Lphn3 and, altogether, to induce excitatory synapse formation. The DHR mutation breaks the interaction of Tenm2 -SS with Lphn3 and abolishes excitatory synapse formation. (B) The Tenm2 +SS isoform does not have rotational flexibility around the linker between the EGF repeats and the rest of the extracellular head as observed in the crystal structure of the Tenm2 +SS isoform, which shows that the splice insert mediates a dimeric interaction between the two Tenm2 +SS protomers (Jackson et al., 2018) (Figure S7, PDB ID: 6FB3). Instead, the Tenm2 +SS protomers are zipped-up due to the additional two salt bridges (black balls) between the propellers of the Tenm2 cis-dimer. Thus, the Lphn-binding site on Tenm2 +SS is not at the right docking geometry to interact with the Lec domain of Lphn3 expressed on the neighboring cell (though it can still bind soluble Lec domain). The geometry of Tenm2 +SS likely enables other hetero- or homophilic protein interactions that were not possible in the -SS isoform, such as Tenm2 trans-homodimerization, and mediates inhibitory synapse formation. The DHR mutation on Tenm2 +SS has no effect on these unknown interactions and thus, does not affect the ability of Tenm2 +SS to induce inhibitory synapses. Model partially drawn to scale. The Lphn ECR structure (orange), is based on Lec and Olf domain structure (PDB: 5AFB), connected by a STP-rich stalk to the GAIN and HormR domain structure (PDB: 4DLQ). Tenm2 protomers are colored as cyan and palegreen (PDB: 6CMX and 6FB3), FLRT protomers are colored as magenta and gray (PDB: 5CMN).

Importantly, the Lphn3-Tenm2 complex structure revealed that the Lphn3-binding site on Tenm2 is away from the alternatively spliced site that is on the Tenm2 propeller (Figure 1,3), raising the question of how a seven-amino acid splice insert within the >2000 amino acid ECR of Tenm2 could dictate Lphn3 binding and synapse specificity without being close to the binding interface. Alternative splicing in the coding region of proteins expands the functional and regulatory capacity of metazoan genomes (Braunschweig et al., 2013; Chen and Manley, 2009; Kalsotra and Cooper, 2011). In addition to Tenm2, numerous proteins such as DSCAMs, protocadherins, neurexins and neuroligins use alternative splicing for diversifying their functions, such as their ability to bind ligands (Aoto et al., 2013; Fuccillo et al., 2015; Irimia et al., 2014; Thalhammer et al., 2017). However, in most proteins, the alternatively spliced sites localize to the ligand-binding site in order to directly enable or disturb ligand binding (Arac et al., 2007; Shen et al., 2008; Wilson et al., 2019). Thus, it is unusual that the Lphn3 binding site is localized away from the alternatively spliced sequence. The only exception we are aware of is a recent study which reported that alternative splicing switches the closed conformation of the large ECR of GPR126, an adhesion GPCR involved in Schwann cell myelination, to a more open and dynamic conformation and consequently increases basal signaling of the receptor (Leon et al., Net Comm., accepter in principle). In the case of Tenm2, alternative splicing allows the protein to act as a switch in regulating ligand binding despite the ligand-binding site being away from the seven residue alternatively spliced site (Li et al., 2018), and this switch disables Lphn3 binding that is required for excitatory synapse formation while likely enabling (an)other interaction(s) required for inhibitory synapse formation.

Although it is intuitively difficult to understand the relationship between alternative splicing and Lphn3 binding, our synapse formation experiments demonstrated a clear requirement of Tenm2-Lphn3 interaction for the excitatory synapse specification function of Tenm2 -SS, since the DHR mutation on Tenm2 -SS isoform that is unable to bind to Lphn3 was unable to induce excitatory postsynaptic specializations (Figure 6B-C). The same mutation on the Tenm2 +SS isoform, however, behaved like wild type Tenm2 +SS and successfully induced inhibitory synapse formation (Figure 6D-E). These results suggest that interaction of Tenm2 with Lphn3 is required for excitatory but not for inhibitory synapse formation. Moreover, the observation that the DHR Lphn-binding mutant had no effect on the ability of Tenm2 +SS to induce inhibitory postsynaptic specializations suggests a Lphn-independent mechanism that requires unidentified Tenm2 interaction partners at inhibitory synapses.

Our results show that the interaction of Lphn3 with Tenm2 can be disrupted in at least two ways: 1) by point mutations on the Lphn-binding interface on Tenm2 (but not by mutations that are not at the interface, Figure S6), and 2) by insertion of the seven alternatively spliced residues in the propeller domain of Tenm2. The mutagenesis of the Lphn-binding interface abolished the Tenm2/Lphn3 interaction in all experimental setups as expected from a binding site mutant (Figure 4). However, the effect of alternatively spliced site on the Tenm2/Lphn3 interaction depended on whether one or both proteins experienced restraints due to their attachment to cell-membranes; or they could freely rotate and tumble in solution (Figure 5). Specifically, alternative splicing abolished the Tenm2/Lphn3 interaction in cell-aggregation experiments where proteins approach each other from opposing membranes; but not in cell-surface staining or in-solution experiments where one or more proteins are in solution. Altogether these results suggest that alternative splicing regulates the Tenm2/Lphn3 interaction via a mechanism that differs from disrupting the binding interface. These results enable us to suggest a speculative model for how alternative splicing regulates Tenm2 interactions and functions:

Tenm2 forms a cis-dimer on the presynaptic membrane that is mediated by two disulfide bonds formed between the 2^nd^ and 5^th^ EGF repeats (black lines, Figure 7) that extend the globular cytoplasmic C-terminal heads of Tenm2 (Tenm2 ECRd1) towards the opposite membrane. Previous cryo-EM images of the dimeric Tenm2 -SS showed that the globular heads have the rotational flexibility around the EGF/head linker (arrow) that enables Tenm2 -SS to sample the 3D space (Li et al., 2018) and to successfully bind the Lec domain of Lphn3 in *cis* and *trans* (Figure 7A). In this conformation, FLRT3 is also able to interact with Lphn3 and form a trimeric complex, consequently leading to excitatory synapse formation (Figure 7A). However, the crystal structure of the Tenm2 +SS isoform showed that, in the presence of the splice insert, the two globular heads form a dimer that is facilitated by the interactions between the splice inserts (Figure S7) (Jackson et al., 2018). The presence of the splice inserts enables the formation of two salt bridges (E1306-H1315 and E1301-R1337 in chicken Tenm2, black balls in Figure 7B, red dashed lines in Figure S7) and five hydrogen bonds between the beta propellers. These newly generated interactions of the propeller domains would zipper-up the molecule introducing rigidity to the Tenm2 +SS cis-dimer and restrict rotational flexibility around the EGF/head linker preventing Tenm2 +SS from sampling the 3D space (Figure 7B). On the other hand, the extracellular domain of Lphn3 consists of two globular regions separated by a Ser-Thr-Pro rich glycosylated linker region that is reported to be semi-rigid (O’Sullivan et al., 2014). As a result, the Lec domain of Lphn3 on the opposite membrane would have limited or no access to the lphn-binding site on Tenm2 +SS and fail to bind, although the binding site is intact and functional. As alternative splicing prevents the formation of the Tenm2/Lphn3 interaction by hiding it rather than destroying it, it is plausible that the +SS isoform enables other Tenm2 interactions that the -SS isoform cannot mediate. Indeed, the trans-dimerization of Tenm2 was reported to occur only by the +SS isoform (Berns et al., 2018); and previous studies suggested that Tenm2 +SS isoform should interact with unknown ligands in order to induce inhibitory synapses (Li et al., 2018). This speculative model explains the currently available data. This first structure of a teneurin-latrophilin complex in combination with our biochemical results demonstrate the clear mechanistic difference of excitatory vs. inhibitory synapse specification and lead to previously unimagined new directions in both the synapse formation and alternative splicing fields and will form the basis for further investigations.

## ACKNOWLEDGEMENTS

We thank Tobin R. Sosnick for generously providing DesG tag construct to improve behavior of the protein, Anthony Kossiakoff, Engin Ozkan for the use of flow cytometer, Georgios Skiniotis and Moran Shalev-Binami for initial data collection. This research was in part, supported by the National Cancer Institute’s National Cryo-EM Facility at the Frederick National Laboratory for Cancer Research under contract HSSN261200800001E”. This work was supported by grants R01 GM120322 (to DA) and R01 GM134035-01 (to DA) and the American Heart Association grant #19POST34380439/Jingxian Li/2019-2020 (to JX).

## AUTHOR CONTRIBUTIONS

J.L. and D.A. designed all experiments and interpreted results. J.L. cloned, expressed and purified Tenm2/Lphn3 complex, carried out Tenm2 related biochemical characterizations and specimen screening, designed and performed the HEK cell expression and flow cytometry binding assays. Y.X. and M.Z. performed cryo-EM data collection and map calculation, model building and refinement. Y.X. and M.Z. carried out structural analysis with assistance from J.L. and D.A. T.C.S., R.S., X.J. and S.C designed and performed the cell aggregation assays, cell surface staining assays and synapse formation assays. S.K participated in Tenm2 alternative splice site related biochemical characterizations. M.P. and K.L assisted in specimen screening by EM. D.A. wrote the paper with assistance from M.Z. T.C.S. J.L. and Y.X. D.A. and M.Z. supervised the project.

## STAR METHODS

### Cell Culture

High-Five insect cells (Trichoplusia ni, female, ovarian) cultured in Insect-Xpress medium (Lonza) supplemented with 10 μg/mL gentamicin at 27°C were used for production of recombinant proteins. HEK293T mammalian cells were used for cell-surface expression assays and flow cytometry binding assays and were cultured in Dulbecco’s modified Eagle’s medium (DMEM; Gibco) supplemented with 10% FBS (Sigma) at 37°C in 5% CO_2._

### Cloning and expression in insect cells

Tenm2 splice variant Lasso (UniProt: Q9NT68-2) and Lphn (Lphn1, UniProt: O88917; Lphn3, UniProt: Q9HAR2) constructs were cloned into a pAcGP67a vector and expressed in High-Five insect cells using the baculovirus expression system as previously described (Arac et al., 2012).

For the structural studies, Tenm2 ECRΔ1 (residues T727-R2648) and LPNH3 ECR (residues S21-V866) were cloned with carboxyl-terminal 6XHis-tags separately and co-expressed in High-Five insect cells. 72 hrs after viral infection, the medium containing secreted glycosylated proteins was collected and the complex was purified with immobilized metal-affinity chromatography (Ni^2+^-NTA agarose resin; QIAGEN) as previously described (Li et al., 2018), followed by size-exclusion chromatography (Superose 6 Increase 10/300 columns, Superdex 200 10/300 GL; GE Healthcare) in 10 mM Tris (pH 8.5), 150 mM NaCl. For the flow cytometry binding assays, Lphn1 Lec (residues S26-Y131) and Lphn3 Lec (residues S21-Y126) were cloned with carboxyl-terminal 6XHis-AVI-tags and purified as described before (Li et al., 2018).

### Cloning and expression in mammalian cells

Full-length Tenm2 (residues M1-R2648) construct and Tenm2 mutants (“DHR” mutant: D1737N, H1738T, R1739T; “LYR” mutant: L1990N, R1992T) bearing HA-tag (inserted between K405/E406) and carboxyl-terminal FLAG-tag were cloned into a pcDNA3.1 vector for cell-surface expression assays and flow cytometry binding assays in HEK293T cells. Tenm2 mutants were generated with a standard two-step PCR-based strategy.

### Flow cytometry

HEK293T cells were cultured in 6-well plates and were transfected 2 μg cDNA using LipoD293 transfection reagent as described before (Li et al., 2018). Transfection was performed with cells at 50-60% confluence. The cells were detached using citric saline solution (50 mM sodium citrate, 135 mM KCl) after 48 hr incubation and washed with PBS + 2% BSA. To test Tenm2 WT and mutant cell surface expression, cells were stained with a primary antibody mixture: mouse anti-FLAG M2 1:1000 and rabbit anti-HA 1:1000 for 30 min at room temperature. After wash with PBS + 2% BSA, cells were stained with a secondary antibody mixture: donkey anti-mouse Alexa Fluor 488 1:3000 and goat anti-rabbit Alexa Fluor 647 1:3000 for 30 min. Cell pellets were resuspended in PBS + 2% BSA immediately before flow cytometry data acquisition (Accuri C6 flow cytometer, 10000 events measured) after washing. Acquired data was analyzed using the FlowJo analysis software (FlowJo LLC).

For the binding assays, His-Avi-tagged Lec was captured on nickel-nitrilotriacetic resin and biotinylated as shown previously (Li et al., 2018). The efficiency of biotinylation was assessed using streptavidin beads. Purified protein was applied to size-exclusion chromatography in 10 mM Tris (pH 8.5), 150 mM NaCl. Biotinylated Lec was tetramerized and fluorescently labeled through incubation with NeutrAvidin DyLight 488 on ice for 20 min. Cultured cells expressing HA-tagged Tenm2 were detached and then washed as described above. Next, the cells were stained with rabbit anti-HA 1:1000 antibody and, following two wash cycles, stained with goat anti-rabbit Alexa Fluor 647 antibody in the presence of the 100 nM NAV488 labeled Lec mixture.

### Cell aggregation assays

HEK293T cells (ATCC) were grown to 90% confluence in a T-75 flask. Cells were trypsinized with 3 mL 0.05% trypsin-EDTA (Gibco Cat#25300-054) and resuspended to 10 mL with DMEM/10% FBS/1% Penicillin-Streptomycin media (Complete DMEM). Three-hundred µL of the cell suspension was added to each well of a 6-well plate containing 3 mL of Complete DMEM media and incubated overnight at 37°C. Cells in each well were then co-transfected with 2 µg of either pCMV (empty vector) + pEmerald, pCMV Lphn3 + pEmerald, pCMV (empty vector) + pCMV dsRed, or dsRed and the indicated Tenm2 construct using the Calcium Phosphate method. All cDNAs were encoded in the pCMV5 or pcDNA3 vector and driven by the CMV promoter. Three days after transfection, the media was aspirated and cells were gently washed with 2 mL of PBS. Cells were resuspended by adding 1 mL of Resuspension Solution (PBS containing 1 mM EGTA) and then incubated for 5 min at 37°C. Fifteen µL of 1 mg/20 µL DNAse (Sigma Cat#D5025) was then added to each well and cells were triturated by pipetting up-and-down (16 times) in each well to resuspend cells off the plate bottom and create single-cell suspensions. Cells were then transferred to a new Eppendorf tube and another 15 µL of DNAse solution was added to each sample. Cells were mixed in 1:1 ratio by adding 70 µL of pCMV (empty vector) + pEmerald or Lphn3 + pEmerald with 70 µL of pCMV (empty vector) + pCMV-dsRed or Tenm2 Construct + dsRed in a new Eppendorf that contained 360 µL of Incubation Solution (DMEM containing 50 mM HEPES-NaOH pH 7.4, 10% FBS, 10 mM CaCl_2_ and 10 mM MgCl_2_) for a final volume of 500 µL. The mixture was triturated and the entire volume was transferred to one well in a non-coated 12-well plate (Costar Cat#3737). Images were taken immediately (time = 0) using a Leica Fluorescent DMIL LED Microscope with a 10x objective. Cells were then placed on a shaking incubator at 125 rpm at 37°C for 20 minutes and imaged again (time = 20). Aggregation index at time = 20 was calculated using ImageJ, measuring the percentage of signal/frame occupied by cells forming complexes of two or more cells relative to the total signal of the frame.

### Cell surface-binding assays

HEK293T cells (ATCC) were grown to 90% confluence in a T-75 flask. Cells were trypsinized with 3 mL 0.05% trypsin-EDTA (Gibco Cat#25300-054) and resuspended to 10 mL of DMEM + 10% FBS + 1% Penicillin-Streptomycin (complete DMEM) media. Fifty µL of cell suspension was added to each well of a 24-well plate that contained a Matrigel-coated coverslip and 1 mL complete DMEM and incubated overnight. Cells were then co-transfected with 1 µg of either empty pCMV, wild-type Teneurin 2 or the indicated mutant Teneurin construct and 1 ug of pEmerald using the Calcium Phosphate method and incubated for 2 days at 37°C. Transfection media was gently removed and 500 µL of chilled DMEM containing 250 µM of purified, Avi-fusion, biotinylated, rat Lphn1 Lec or human Lphn3 Lec was added to each well. Plates were wrapped in foil and incubated overnight at 4°C to reduce endocytosis, with gentle shaking. This was performed essentially as described in (Boucard et al., 2012). Cells were gently washed 2x using 1 mL of PBS and fixed with 300 µL of ice-cold 4% PFA/4%sucrose/PBS. Plates were wrapped in foil and incubated for 20 minutes at 4°C during the fixation. Cells were gently washed 3x using 1 mL of room temperature PBS and blocked with 300 µL of 5% BSA (Sigma Cat#10735086001)/PBS (blocking buffer) for 1 hr at room temperature. Bound biotinylated Lec was detected by immunofluorescence using 300 µL per well of Streptavidin (AlexaFluor-555 conjugated, Invitrogen Cat#S21381, at 1:10,000 dilution) diluted into blocking buffer for 1 hr at room temperature. Cells were gently washed 3x with 1 mL of PBS. Cells were re-blocked, and HA-tagged, surface Teneurins were detected by adding 300 µL of rabbit anti-HA antibody (Cell Signaling Technologies Cat#3724) at 1:1,000 dilution in blocking buffer. Cells were gently washed 3x with 1 mL PBS. Goat anti-rabbit secondary antibodies (Alexa-Fluor 633 conjugated, Invitrogen) and DAPI (Sigma Cat#10236276001) staining was done for 30 minutes at 1:10,000 and 1:5,000, respectively, in blocking buffer, followed by 3x gentle washes with 1 mL of PBS. Coverslips were mounted onto slides (UltraClear microscope slides Denville Scientific Cat# M1021) in mounting media (Fluoromount-G, Southern Biotech Cat#010020). Images were acquired using a Nikon A1 Eclipse Ti2 confocal microscope with a 60x oil-immersion objective, operated by NIS-Elements AR acquisition software. The same confocal acquisition settings were applied to all samples of the experiment. Collected z-stacks at a 0.4 µm z-step size were analyzed blindly using Nikon Elements Analysis software. Co-localization was calculated using the Pearson’s correlation coefficient of Lec-Streptavidin-555 to Teneurin-HA-633 emission.

### Artificial synapse formation assay

HEK293T cells were transfected with the expression vectors of the cell adhesion molecules. 24 hrs later, HEK293T cells were co-cultured with cultured cortical neurons (DIV16) from P0 mice. After 24 hrs, cells were fixed with 4% PFA and immunostained with rabbit anti-Flag (Sigma; 1:1000 both) together with mouse anti-PSD95 (Sysy; 1:500) or mouse anti-GABAA α2 (Sysy; 1:500) respectively. Images were collected with a Nikon A1 confocal microscope using a 60x objective. A human NPR mutant (1-118 aa of the full-length protein) which comprises an A domain containing low-complexity sequences is used as the negative control in the artificial synapse formation assays. The signals of the synaptic markers that were recruited to the surface of the HEK293T cells were quantified using Image J. Normalized values equal the fluorescent intensity of the synaptic marker that was examined (GABAγ2/PSD95) / the fluorescent intensity of the Flag-tagged protein expressed in the HEK cells (Tenm’s /Nrn1β).

### Cryo-EM data acquisition

2.5μl purified human Tenm2 ECRΔ1 and human Lphn3 ECR complex (0.22□mg/ml) was applied on glow-discharged holey carbon grids (Quantifoil R1.2/1.3, 300 mesh), and vitrified using a Vitrobot Mark IV (FEI Company). The specimen was visualized using a Titan Krios electron microscope (FEI) operating at 300□kV and equipped with a K3 direct electron detector (Gatan, Inc.). Images were recorded with a nominal magnification of 81,000x in super-resolution counting mode, corresponding to a pixel size of 0.54□Å on the specimen level. To maximize data collection speed while keeping image aberrations minimal, image shift was used as imaging strategy using one data position per hole with four holes targeted in one template with one focus position. In total, 4,967 images with defocus values in the range of −1.0 to −2.5 μm were recorded using a dose rate of 14.6 electrons/Å2/s. The total exposure time was set to 4.2 s with frames recorded every 0.105 s, resulting in an accumulated dose of about 60.1 electrons per Å2 and a total of 40 frames per movie stack.

### Image processing and 3D reconstructions

Stack images were subjected to beam-induced motion correction using MotionCor2 (Glodeck et al., 2016). CTF parameters for each micrograph were determined by CTFFIND4 (Mindell and Grigorieff, 2003). Particle selection, two- and three-dimensional classifications were initially performed on a binned dataset with a pixel size of 4.32□Å using RELION-3 (Zivanov et al., 2018). In total, 4,475,958 particle projections were selected using automated particle picking and subjected to reference-free two-dimensional classification to discard false-positive particles or particles categorized in poorly defined classes, resulting in 3,307,148 particle projections for further processing. The initial 3D maximum-likelihood-based classification was performed on a binned dataset with a pixel size of 4.32□Å using the previously reported Tenm2 structure (Li et al., 2018) as the reference model. The detailed data processing flow is shown in Fig. S2 and S3. Briefly, for Tenurin-Letrophilin complex, 1,309,684 particles that showed well-defined density of Lec domain were selected after initial rounds of 3D classification. Then, two rounds of focused 3D classification with mask around Lec domain were performed without alignment. 3D refinement and post-processing was performed on the best class with clear features for the Lec domain (Fig. S2). The final map for Tenm2_Lec was resolved at 2.97Å (Fig. S2). To resolve domain 3, the same dataset was reprocessed with a total of 1,137,765 particles showing well-resolved domain3. Two rounds of focused 3D classification with mask around domain3 were performed, followed by 3D refinement and post-processing. The final map for Tenm2_domain3 was resolved at 3.07Å (Fig. S3).

Reported resolutions are based on the gold-standard Fourier shell correlation (FSC) using the 0.143 criterion (Fig. S1C). All density maps were corrected for the modulation transfer function (MTF) of the K3 direct detector and then sharpened by applying a temperature factor that was estimated using post-processing in RELION-3. Local resolution was determined using ResMap (Kucukelbir et al., 2014) with half-reconstructions as input maps (Fig. S1D).

### Model building and refinement

Model building was based on the structure of human Tenm2 −2 extracellular region (PDB: 6CMX) and the Lec domain from human Lphn3 −3 (PDB: 5AFB and 5FTT). The models were first docked into the EM density maps using Chimera (Pettersen et al., 2004) and then manually adjusted residue-by-residue to fit the density using COOT (Emsley et al., 2010). The ECR of Tenm2 was built based on that of chicken Tenm2 (PDB: 6FB3) and manually adjusted to human sequence and splice form. Both maps (Ten_Lec and Ten_domain3) were used for model building. There is a slight shift between the two maps from reconstruction, so they were aligned based on Ten-Lec before model building. The final model containing both ECR of Tenm2 −2 and Lec domain of Letrophilin-3 was subjected to global refinement and minimization in real space using the *phenix_real_space_refine* module in Phenix (Adams et al., 2010) first against Ten_domain3 map while keeping the Lec domain as a rigid body, and then against Ten_Lec map while keeping the ECR3 as a rigid body. FSC curves were calculated between the resulting model and either maps using Phenix M-triage (Figure S2). The final model statistics are provided in Table 1.

### Quantification and Statistical Analysis

Error bars in Figure 4, 5, 6 and Figure S6 represent means ± SEM. Each measurement was repeated at least three times independently. Data was analyzed using software GraphPad Prism and ImageJ.

## Data and Software Availability

### Data Resources

The cryo-EM density map has been deposited in the Electron Microscopy Data Bank (https://www.ebi.ac.uk/pdbe/emdb/) under accession code EMD-XXXX and the model coordinates have been deposited in the Protein Data Bank (http://www.rcsb.org) under accession number XXXX.

## SUPPLEMENTARY FIGURE LEGENDS

**Figure S1. Map Quality and Local Resolution, related to Figure 1**

(A) A representative electron micrograph of TEN/Lphn3 complex collected using a K3 direct detection camera (Gatan, Inc., Pleasanton, CA). Scale bar is 50 nM. (B) Representative 2D class averages of TEN/Lphn3 complex. The diameter of the circular mask is 180 Å. The Lec domain is highlighted in yellow circles. (C) Fourier shell correlation (FSC) curves of the 3D density maps after RELION post-processing. The resolutions are determined using FSC=0.143 criterion. (D) Two views of the final 3D density maps of TEN_Lphn3 complex and TEN-Domain3 are colored based on the local resolution determined by ResMap (Kucukelbir et al., 2014). Corresponding angular distribution plots for all particles in the final maps were shown under each view.

**Figure S2. Single-particle cryo-EM image processing workflow for the TEN_Lphn3 complex, related to Figure 1**

The single-particle cryo-EM dataset for TEN/Lphn3 complexes were subjected to particle selection, 2D classification, and rounds of 3D classification using a low-pass filtered model of Tenm2 (Li et al., Cell 2018) as the initial model. Classes with clear densities of the Lec domain (highlighted in red circles) were further selected for focused 3D classification with a mask surrounding the Lec domain. The best class was selected for 3D refinement and post-processing, resulting in the final map of TEN-Lphn3 complex at a nominal resolution of 2.97Å as determined by FSC=0.143 criterion (Fig. S1). Details are provided in the methods section.

**Figure S3. Single-particle cryo-EM image processing workflow for the Tenm2 ECR focusing on domain 3. Related to Figure 1**

After initial rounds of 3D classification using a low-pass filtered model of Tenm2 (Li et al., Cell 2018) as the initial model (Fig. S2), classes with clear densities of Tenm2 domain 3 (highlighted in red circles) were further selected for a focused 3D classification with a mask surrounding domain 3. The best class was selected for 3D refinement and post-processing, resulting in a final map of Tenm2 with well-resolved domain 3 at a nominal resolution of 3.07Å as determined by FSC=0.143 criteria (Fig. S1). Details are provided in the methods section.

**Figure S4. Model quality, Related to Figure 1**

Snapshots of map versus model agreement. Each region is highlighted by a dashed box including the representative regions of an α-helix in Tenm2 Tox-like domain (A), a β-sheet in Tenm2 β-barrel domain (B), the Lec domain of Lphn3 (C), and the entire Tenm2 β-propeller domain (D). The alternative splice site in β-propeller domain is located between Arginine 1156 and Asparagine 1157 (highlighted in pink). Contour level for individual snapshots: A, 5.0 rmsd; B, 5.0 rmsd; C, 4.0 rmsd; D, 5.0 rmsd.

**Figure S5. Conservation, glycosylation and mutagenesis of the Lphn-binding site on Tenm2, Related to Figure 3**

(A) Conservation analysis of Tenm2 shows a highly conserved patch on the barrel surface (indicated by yellow circles) opposite to the Tox-like domain. This conserved patch is not present in bacterial toxins, implying the evolutionary importance of this region in Tenm2, which may be employed to interact with partner proteins. (B) Comparison of potential sugar binding site of Lphn3 Lec domain with other SUEL-related Lec family members. Lec domain in human Lphn3 with density of glycochain from human Tenm2 shown in grey mesh is colored as orange; Lec domain in mouse Lphn1 from NMR model (PDB: 2JXA) with rhamnose highlighted in red is colored as yellow (Vakonakis et al., 2008). Three Lec repeats in sea urchin Lphn3 (PDB: 5H4S) with rhamnose highlighted in red are colored as pink, palecyan and blue, respectively (Hatakeyama et al., 2017). All the Lec domains are aligned. Note that the glycochain density from Tenm2 is close to the sugar binding pocket of other Lec domains.

(C) Flow cytometry analysis of cell-surface expression for full length Tenm2 on non-permeabilized HEK293T mammalian cells, compared to empty vector-transfected cells (EV) and cells that were transfected with mutant Tenm2 constructs.

**Figure S6. LR mutation on Tenm2 has no influence on Lphn3 binding; Toxin-like domain deletion results, Related to Figure 3**

(A) Diagram for WT Tenm2 +SS, Tenm2 LR +SS (L1990N, R1992T), Tenm2 +SS Tox domain deletion (ΔTox) and Tenm2 +SS barrel and Tox domain deletion (ΔTox ΔBarrel) constructs that were used in this and our previous study (Li et al., 2018). Tenm2 LR +SS construct is generated as a positive control and it carries two point mutations in the Tenm2 barrel which are not at the lphn-binding site. The Tenm2 +SS ΔTox and Tenm2 +SS ΔTox ΔBarrel are the same constructs as was used in Li et al.

(B) Cell surface staining assays suggest that the “LR” mutations on Tenm2 β-barrel has no influence on the binding between Tenm2 and Lphn3. Cell surface staining assays suggest that the toxin-domain deletion abolishes Lphn binding.

Supplementary Discussion: In Li et al., we reported that the toxin-like domain is needed for Lphn3 binding because the Tenm2 +SS ΔTox that deletes the toxin domain of Tenm2 had abolished Lec binding in cell-aggregation and cell-surface staining experiments and lacked any defects in cell-surface localization. These results mislead us to pinpoint the exact binding interface for Lec. We have repeated the experiments and confirmed these results are reproducible. We believe that this mutant escapes the protein quality control system and is localized on the cell surface although it is misfolded and thus is unable to bind Lphn. However, the real Lphn-binding site is not the toxin-like domain.

Scale bar indicates 20 um. Quantification of cell surface binding assays are shown next to the images.

(C) Wild-type (WT) and mutant Tenm2 proteins were tested for surface expression in HEK293T cells as well as their ability to bind soluble Lphn1/Lphn3 Lec domain using flow cytometry. Data relate to Tenm2 +SS delete Tox domain (ΔTox) and Tenm2 +SS delete Barrel and Tox domain (ΔTox Δ Barrel) are modified from Li et al., 2018 and separated by black line.

**Figure S7. Alternatively-spliced insert within the** β**-propeller mediates the Tenm2 +SS dimer interface, Related to Figure 7**

Structure of Tenm2 +SS dimer shows the splice inserts from each protomer (yellow and magenta residues) creates a binding interface and leads to Tenm2 dimerization via the beta-propeller. Close-up views of the dimer interface show two salt bridges and five hydrogen bonds are at the interface. One of the salt bridges is directly mediated by the glutamate (E1306) within the splice insert NKEFKHS and four of the hydrogen bonds also require splice site residues. Two salt bridges align almost parallel to each other and to the disulfide bonds between the EGF repeats and restricts the conformational flexibility of the Tenm2 head significantly (see Figure 7). The N-termini of both protomers face the same direction towards the EGF repeats, and thus, the dimer is positioned as a cis-dimer that will extend the zippering of the already existing EGF-mediated cis-dimer, although it was reported to form as a trans-homodimer, previously (Jackson et al., 2018). TEN protomers (PDB: 6FB3) are colored as cyan and palegreen, respectively, and splice sites are colored as tellow and magenta, respectively.

## REFERENCES

Aldahmesh, M.A., Mohammed, J.Y., Al-Hazzaa, S., and Alkuraya, F.S. (2012). Homozygous null mutation in ODZ3 causes microphthalmia in humans. Genet Med 14, 900–904.

Alkelai, A., Olender, T., Haffner-Krausz, R., Tsoory, M.M., Boyko, V., Tatarskyy, P., Gross-Isseroff, R., Milgrom, R., Shushan, S., Blau, I., et al. (2016). A role for TENM1 mutations in congenital general anosmia. Clin Genet 90, 211–219.

Anderson, G.R., Maxeiner, S., Sando, R., Tsetsenis, T., Malenka, R.C., and Sudhof, T.C. (2017). Postsynaptic adhesion GPCR latrophilin-2 mediates target recognition in entorhinal-hippocampal synapse assembly. The Journal of cell biology 216, 3831–3846.

Aoto, J., Martinelli, D.C., Malenka, R.C., Tabuchi, K., and Südhof, T.C. (2013). Presynaptic neurexin-3 alternative splicing trans-synaptically controls postsynaptic AMPA receptor trafficking. Cell 154, 75–88.

Arac, D., Boucard, A.A., Ozkan, E., Strop, P., Newell, E., Sudhof, T.C., and Brunger, A.T. (2007). Structures of neuroligin-1 and the neuroligin-1/neurexin-1 beta complex reveal specific protein-protein and protein-Ca2+ interactions. Neuron 56, 992–1003.

Arcos-Burgos, M., Jain, M., Acosta, M.T., Shively, S., Stanescu, H., Wallis, D., Domene, S., Velez, J.I., Karkera, J.D., Balog, J., et al. (2010). A common variant of the latrophilin 3 gene, LPHN3, confers susceptibility to ADHD and predicts effectiveness of stimulant medication. Molecular psychiatry 15, 1053–1066.

Baumgartner, S., and Chiquet-Ehrismann, R. (1993). Tena, a Drosophila gene related to tenascin, shows selective transcript localization. Mechanisms of development 40, 165–176.

Baumgartner, S., Martin, D., Hagios, C., and Chiquet-Ehrismann, R. (1994). Tenm, a Drosophila gene related to tenascin, is a new pair-rule gene. The EMBO journal 13, 3728–3740.

Berns, D.S., DeNardo, L.A., Pederick, D.T., and Luo, L. (2018). Teneurin-3 controls topographic circuit assembly in the hippocampus. Nature 554, 328–333.

Boucard, A.A., Ko, J., and Sudhof, T.C. (2012). High affinity neurexin binding to cell adhesion G-protein-coupled receptor CIRL1/latrophilin-1 produces an intercellular adhesion complex. The Journal of biological chemistry 287, 9399–9413.

Boucard, A.A., Maxeiner, S., and Sudhof, T.C. (2014). Latrophilins function as heterophilic cell-adhesion molecules by binding to teneurins: regulation by alternative splicing. J Biol Chem 289, 387–402.

Chassaing, N., Ragge, N., Plaisancie, J., Patat, O., Genevieve, D., Rivier, F., Malrieu-Eliaou, C., Hamel, C., Kaplan, J., and Calvas, P. (2016). Confirmation of TENM3 involvement in autosomal recessive colobomatous microphthalmia. Am J Med Genet A 170, 1895–1898.

Deak, F., Liu, X., Khvotchev, M., Li, G., Kavalali, E.T., Sugita, S., and Sudhof, T.C. (2009). Alpha-latrotoxin stimulates a novel pathway of Ca2+-dependent synaptic exocytosis independent of the classical synaptic fusion machinery. The Journal of neuroscience: the official journal of the Society for Neuroscience 29, 8639–8648.

Doyle, S.E., Scholz, M.J., Greer, K.A., Hubbard, A.D., Darnell, D.K., Antin, P.B., Klewer, S.E., and Runyan, R.B. (2006). Latrophilin-2 is a novel component of the epithelial-mesenchymal transition within the atrioventricular canal of the embryonic chicken heart. Dev Dyn 235, 3213–3221.

Feng, K., Zhou, X.H., Oohashi, T., Morgelin, M., Lustig, A., Hirakawa, S., Ninomiya, Y., Engel, J., Rauch, U., and Fassler, R. (2002). All four members of the Ten-m/Odz family of transmembrane proteins form dimers. The Journal of biological chemistry 277, 26128–26135.

Fuccillo, M.V., Foldy, C., Gokce, O., Rothwell, P.E., Sun, G.L., Malenka, R.C., and Sudhof, T.C. (2015). Single-Cell mRNA Profiling Reveals Cell-Type-Specific Expression of Neurexin Isoforms. Neuron 87, 326–340.

Glodeck, D., Hesser, J., and Zheng, L. (2016). Distortion correction of EPI data using multimodal nonrigid registration with an anisotropic regularization. Magn Reson Imaging 34, 127–136.

Graumann, R., Di Capua, G.A., Oyarzun, J.E., Vasquez, M.A., Liao, C., Branes, J.A., Roa, I., Casanello, P., Corvalan, A.H., Owen, G.I., et al. (2017). Expression of teneurins is associated with tumor differentiation and patient survival in ovarian cancer. PloS one 12, e0177244.

Hamann, J., Aust, G., Arac, D., Engel, F.B., Formstone, C., Fredriksson, R., Hall, R.A., Harty, B.L., Kirchhoff, C., Knapp, B., et al. (2015). International Union of Basic and Clinical Pharmacology. XCIV. Adhesion G protein-coupled receptors. Pharmacological reviews 67, 338–367.

Hatakeyama, T., Ichise, A., Unno, H., Goda, S., Oda, T., Tateno, H., Hirabayashi, J., Sakai, H., and Nakagawa, H. (2017). Carbohydrate recognition by the rhamnose-binding lectin SUL-I with a novel three-domain structure isolated from the venom of globiferous pedicellariae of the flower sea urchin Toxopneustes pileolus. Protein science: a publication of the Protein Society 26, 1574–1583.

Hong, W., Mosca, T.J., and Luo, L. (2012). Teneurins instruct synaptic partner matching in an olfactory map. Nature 484, 201–207.

Hor, H., Francescatto, L., Bartesaghi, L., Ortega-Cubero, S., Kousi, M., Lorenzo-Betancor, O., Jimenez-Jimenez, F.J., Gironell, A., Clarimon, J., Drechsel, O., et al. (2015). Missense mutations in TENM4, a regulator of axon guidance and central myelination, cause essential tremor. Human molecular genetics 24, 5677–5686.

Irimia, M., Weatheritt, R.J., Ellis, J.D., Parikshak, N.N., Gonatopoulos-Pournatzis, T., Babor, M., Quesnel-Vallieres, M., Tapial, J., Raj, B., O’Hanlon, D., et al. (2014). A highly conserved program of neuronal microexons is misregulated in autistic brains. Cell 159, 1511–1523.

Jackson, V.A., Meijer, D.H., Carrasquero, M., van Bezouwen, L.S., Lowe, E.D., Kleanthous, C., Janssen, B.J.C., and Seiradake, E. (2018). Structures of Teneurin adhesion receptors reveal an ancient fold for cell-cell interaction. Nature communications 9, 1079.

Kan, Z., Jaiswal, B.S., Stinson, J., Janakiraman, V., Bhatt, D., Stern, H.M., Yue, P., Haverty, P.M., Bourgon, R., Zheng, J., et al. (2010). Diverse somatic mutation patterns and pathway alterations in human cancers. Nature 466, 869–873.

Krasnoperov, V.G., Beavis, R., Chepurny, O.G., Little, A.R., Plotnikov, A.N., and Petrenko, A.G. (1996). The calcium-independent receptor of alpha-latrotoxin is not a neurexin. Biochemical and biophysical research communications 227, 868–875.

Krasnoperov, V.G., Bittner, M.A., Beavis, R., Kuang, Y., Salnikow, K.V., Chepurny, O.G., Little, A.R., Plotnikov, A.N., Wu, D., Holz, R.W., et al. (1997). alpha-Latrotoxin stimulates exocytosis by the interaction with a neuronal G-protein-coupled receptor. Neuron 18, 925–937.

Kucukelbir, A., Sigworth, F.J., and Tagare, H.D. (2014). Quantifying the local resolution of cryo-EM density maps. Nat Methods 11, 63–65.

Lelianova, V.G., Davletov, B.A., Sterling, A., Rahman, M.A., Grishin, E.V., Totty, N.F., and Ushkaryov, Y.A. (1997). Alpha-latrotoxin receptor, latrophilin, is a novel member of the secretin family of G protein-coupled receptors. The Journal of biological chemistry 272, 21504–21508.

Levine, A., Bashan-Ahrend, A., Budai-Hadrian, O., Gartenberg, D., Menasherow, S., and Wides, R. (1994). Odd Oz: a novel Drosophila pair rule gene. Cell 77, 587–598.

Li, J., Shalev-Benami, M., Sando, R., Jiang, X., Kibrom, A., Wang, J., Leon, K., Katanski, C., Nazarko, O., Lu, Y.C., et al. (2018). Structural Basis for Teneurin Function in Circuit-Wiring: A Toxin Motif at the Synapse. Cell 173, 735–748 e715.

Meusch, D., Gatsogiannis, C., Efremov, R.G., Lang, A.E., Hofnagel, O., Vetter, I.R., Aktories, K., and Raunser, S. (2014). Mechanism of Tc toxin action revealed in molecular detail. Nature 508, 61–65.

Mindell, J.A., and Grigorieff, N. (2003). Accurate determination of local defocus and specimen tilt in electron microscopy. J Struct Biol 142, 334–347.

Mosca, T.J., Hong, W., Dani, V.S., Favaloro, V., and Luo, L. (2012). Trans-synaptic Teneurin signalling in neuromuscular synapse organization and target choice. Nature 484, 237–241.

Mosca, T.J., and Luo, L. (2014). Synaptic organization of the Drosophila antennal lobe and its regulation by the Teneurins. eLife 3, e03726.

O’Hayre, M., Vazquez-Prado, J., Kufareva, I., Stawiski, E.W., Handel, T.M., Seshagiri, S., and Gutkind, J.S. (2013). The emerging mutational landscape of G proteins and G-protein-coupled receptors in cancer. Nature reviews Cancer 13, 412–424.

O’Sullivan, M.L., de Wit, J., Savas, J.N., Comoletti, D., Otto-Hitt, S., Yates, J.R., 3rd, and Ghosh, A. (2012). FLRT proteins are endogenous latrophilin ligands and regulate excitatory synapse development. Neuron 73, 903–910.

O’Sullivan, M.L., Martini, F., von Daake, S., Comoletti, D., and Ghosh, A. (2014). LPHN3, a presynaptic adhesion-GPCR implicated in ADHD, regulates the strength of neocortical layer 2/3 synaptic input to layer 5. Neural development 9, 7.

Oohashi, T., Zhou, X.H., Feng, K., Richter, B., Morgelin, M., Perez, M.T., Su, W.D., Chiquet-Ehrismann, R., Rauch, U., and Fassler, R. (1999). Mouse ten-m/Odz is a new family of dimeric type II transmembrane proteins expressed in many tissues. The Journal of cell biology 145, 563–577.

Sando, R., Jiang, X., and Sudhof, T.C. (2019). Latrophilin GPCRs direct synapse specificity by coincident binding of FLRTs and teneurins. Science 363.

Shen, K.C., Kuczynska, D.A., Wu, I.J., Murray, B.H., Sheckler, L.R., and Rudenko, G. (2008). Regulation of neurexin 1beta tertiary structure and ligand binding through alternative splicing. Structure 16, 422–431.

Silva, J.P., Lelianova, V.G., Ermolyuk, Y.S., Vysokov, N., Hitchen, P.G., Berninghausen, O., Rahman, M.A., Zangrandi, A., Fidalgo, S., Tonevitsky, A.G., et al. (2011). Latrophilin 1 and its endogenous ligand Lasso/teneurin-2 form a high-affinity transsynaptic receptor pair with signaling capabilities. Proceedings of the National Academy of Sciences of the United States of America 108, 12113–12118.

Sudhof, T.C. (2001). alpha-Latrotoxin and its receptors: neurexins and CIRL/latrophilins. Annual review of neuroscience 24, 933–962.

Sugita, S., Ichtchenko, K., Khvotchev, M., and Sudhof, T.C. (1998). alpha-Latrotoxin receptor CIRL/latrophilin 1 (CL1) defines an unusual family of ubiquitous G-protein-linked receptors. G-protein coupling not required for triggering exocytosis. The Journal of biological chemistry 273, 32715–32724.

Sugita, S., Khvochtev, M., and Sudhof, T.C. (1999). Neurexins are functional alpha-latrotoxin receptors. Neuron 22, 489–496.

Talamillo, A., Grande, L., Ruiz-Ontanon, P., Velasquez, C., Mollinedo, P., Torices, S., Sanchez-Gomez, P., Aznar, A., Esparis-Ogando, A., Lopez-Lopez, C., et al. (2017). ODZ1 allows glioblastoma to sustain invasiveness through a Myc-dependent transcriptional upregulation of RhoA. Oncogene 36, 1733–1744.

Thalhammer, A., Contestabile, A., Ermolyuk, Y.S., Ng, T., Volynski, K.E., Soong, T.W., Goda, Y., and Cingolani, L.A. (2017). Alternative Splicing of P/Q-Type Ca(2+) Channels Shapes Presynaptic Plasticity. Cell reports 20, 333–343.

Vakonakis, I., Langenhan, T., Promel, S., Russ, A., and Campbell, I.D. (2008). Solution structure and sugar-binding mechanism of mouse latrophilin-1 RBL: a 7TM receptor-attached lectin-like domain. Structure 16, 944–953.

Vysokov, N.V., Silva, J.P., Lelianova, V.G., Ho, C., Djamgoz, M.B., Tonevitsky, A.G., and Ushkaryov, Y.A. (2016). The Mechanism of Regulated Release of Lasso/Teneurin-2. Frontiers in molecular neuroscience 9, 59.

Wilson, S.C., White, K.I., Zhou, Q., Pfuetzner, R.A., Choi, U.B., Sudhof, T.C., and Brunger, A.T. (2019). Structures of neurexophilin-neurexin complexes reveal a regulatory mechanism of alternative splicing. The EMBO journal 38, e101603.

Woelfle, R., D’Aquila, A.L., and Lovejoy, D.A. (2016). Teneurins, TCAP, and latrophilins: roles in the etiology of mood disorders. Transl Neurosci 7, 17–23.

Ziegler, A., Corvalan, A., Roa, I., Branes, J.A., and Wollscheid, B. (2012). Teneurin protein family: an emerging role in human tumorigenesis and drug resistance. Cancer Lett 326, 1–7.

Zivanov, J., Nakane, T., Forsberg, B.O., Kimanius, D., Hagen, W.J., Lindahl, E., and Scheres, S.H. (2018). New tools for automated high-resolution cryo-EM structure determination in RELION-3. eLife 7.

