## Supplementary information for "Structure of the teneurin-latrophilin complex: Alternative splicing controls synapse specificity by a novel mechanism"

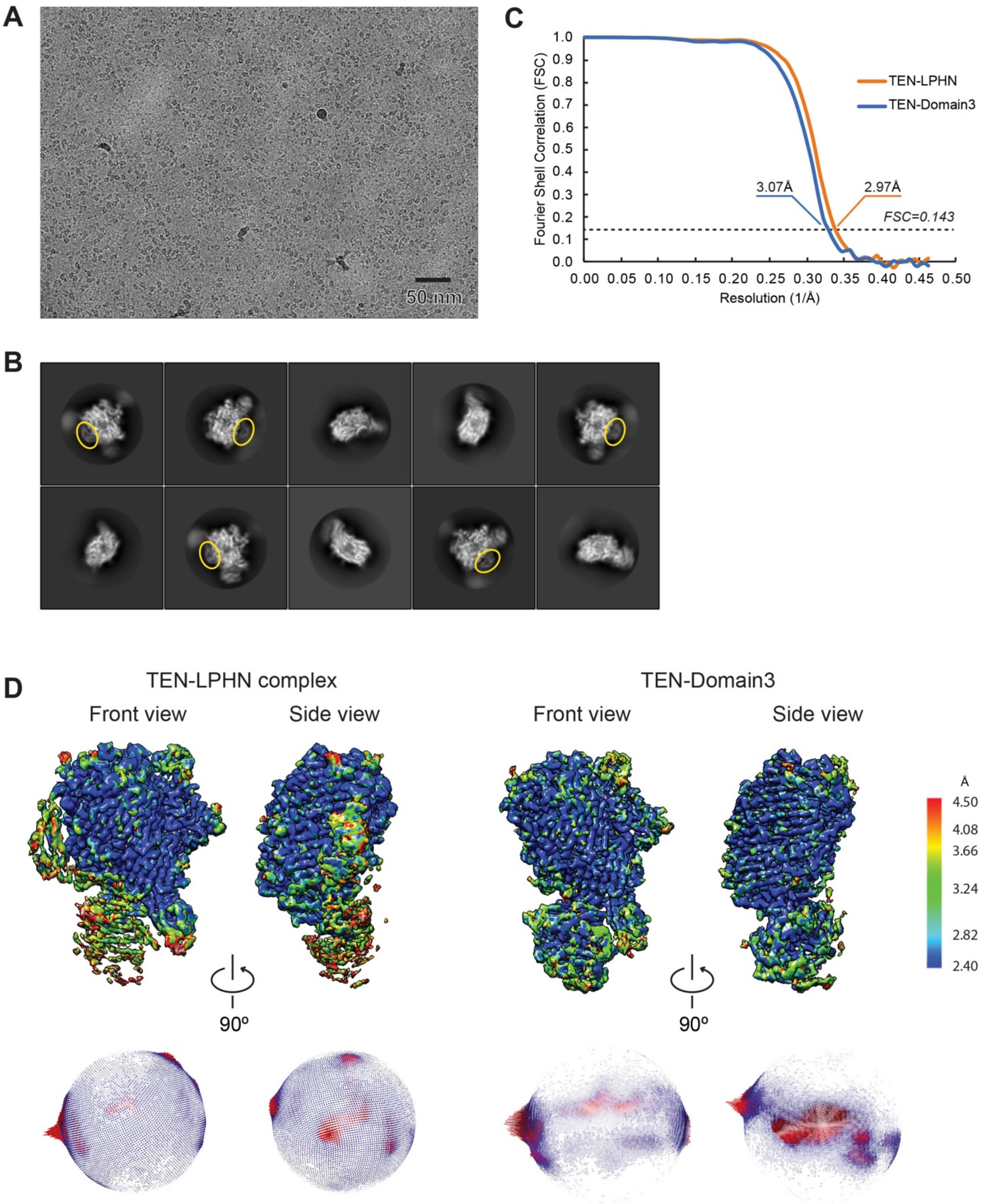


**Figure S1. Map Quality and Local Resolution, related to Figure 1**

(A) A representative electron micrograph of TEN/Lphn3 complex collected using a K3 direct detection camera (Gatan, Inc., Pleasanton, CA). Scale bar is 50 nM. (B) Representative 2D class averages of TEN/Lphn3 complex. The diameter of the circular mask is 180 Å. The Lec domain is highlighted in yellow circles. (C) Fourier shell correlation (FSC) curves of the 3D density maps after RELION post-processing. The resolutions are determined using FSC=0.143 criterion. (D) Two views of the final 3D density maps of TEN_Lphn3 complex and TEN-Domain3 are colored based on the local resolution determined by ResMap (Kucukelbir et al., 2014). Corresponding angular distribution plots for all particles in the final maps were shown under each view.

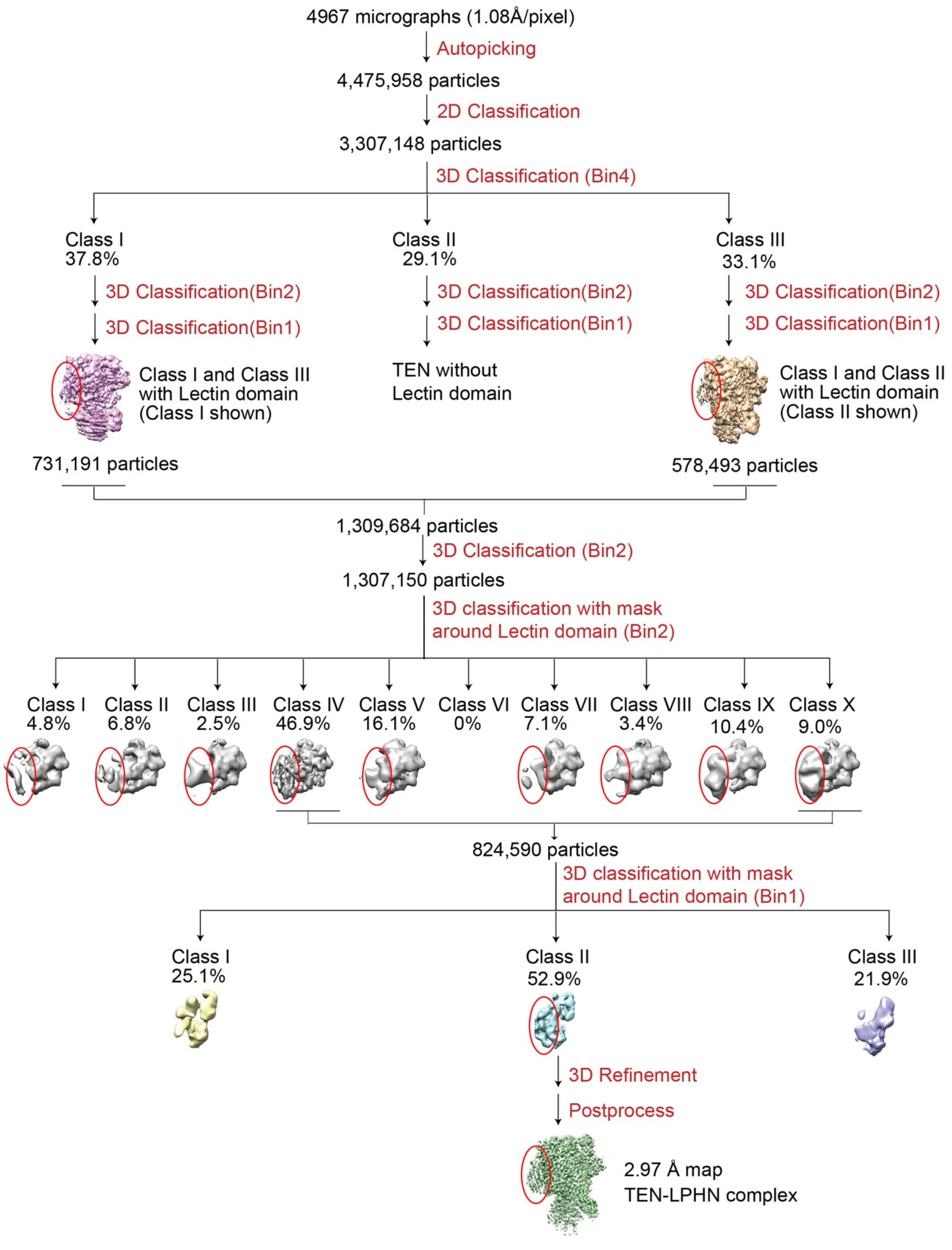


**Figure S2. Single-particle cryo-EM image processing workflow for the TEN_Lphn3 complex, related to Figure 1**

The single-particle cryo-EM dataset for TEN/Lphn3 complexes were subjected to particle selection, 2D classification, and rounds of 3D classification using a low-pass filtered model of Tenm2 (Li et al., Cell 2018) as the initial model. Classes with clear densities of the Lec domain (highlighted in red circles) were further selected for focused 3D classification with a mask surrounding the Lec domain. The best class was selected for 3D refinement and post-processing, resulting in the final map of TEN-Lphn3 complex at a nominal resolution of 2.97Å as determined by FSC=0.143 criterion (Fig. S1). Details are provided in the methods section.

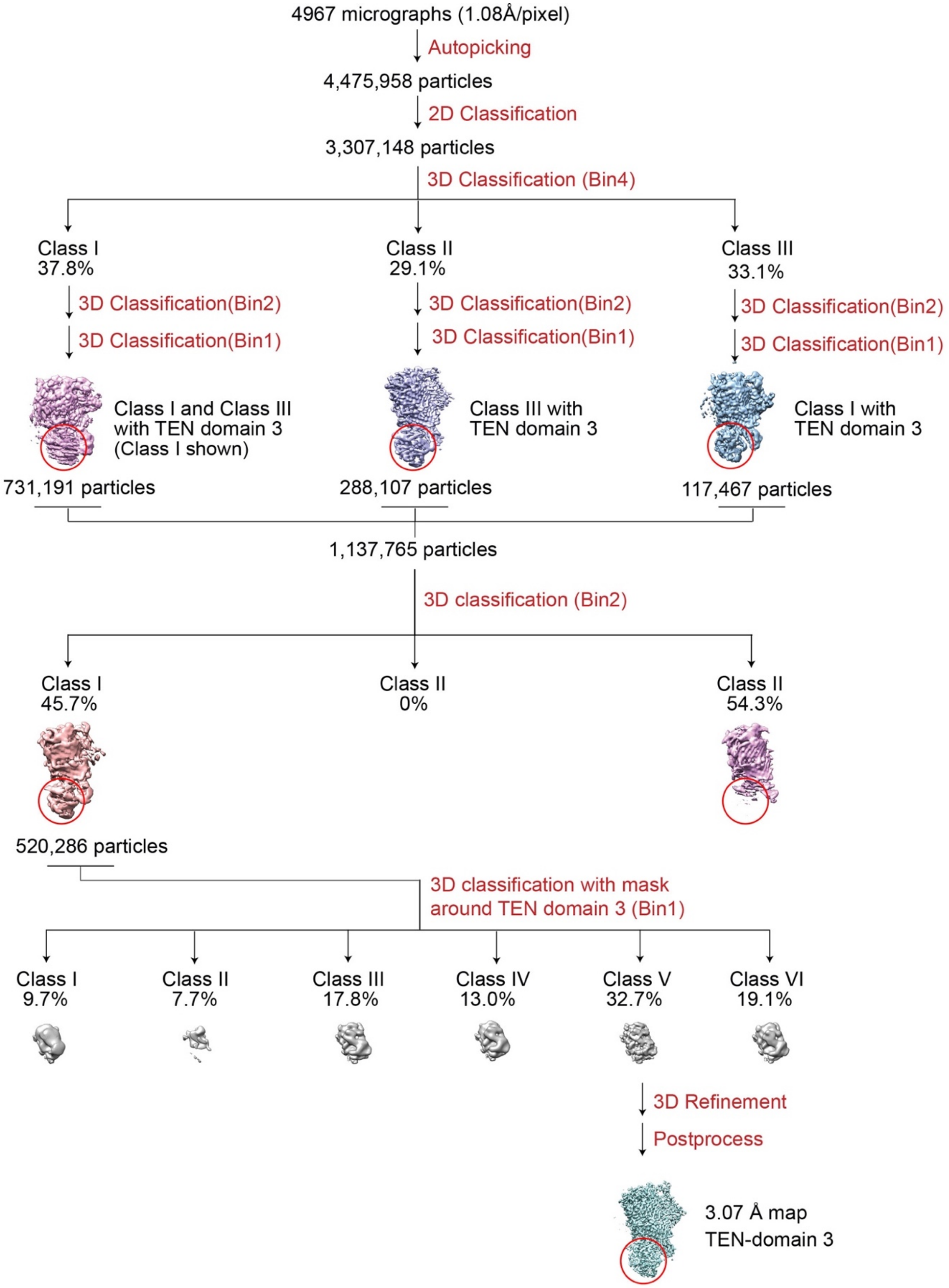


**Figure S3. Single-particle cryo-EM image processing workflow for the Tenm2 ECR focusing on domain 3. Related to Figure 1**

After initial rounds of 3D classification using a low-pass filtered model of Tenm2 (Li et al., Cell 2018) as the initial model (Fig. S2), classes with clear densities of Tenm2 domain 3 (highlighted in red circles) were further selected for a focused 3D classification with a mask surrounding domain 3. The best class was selected for 3D refinement and post-processing, resulting in a final map of Tenm2 with well-resolved domain 3 at a nominal resolution of 3.07Å as determined by FSC=0.143 criteria (Fig. S1). Details are provided in the methods section.

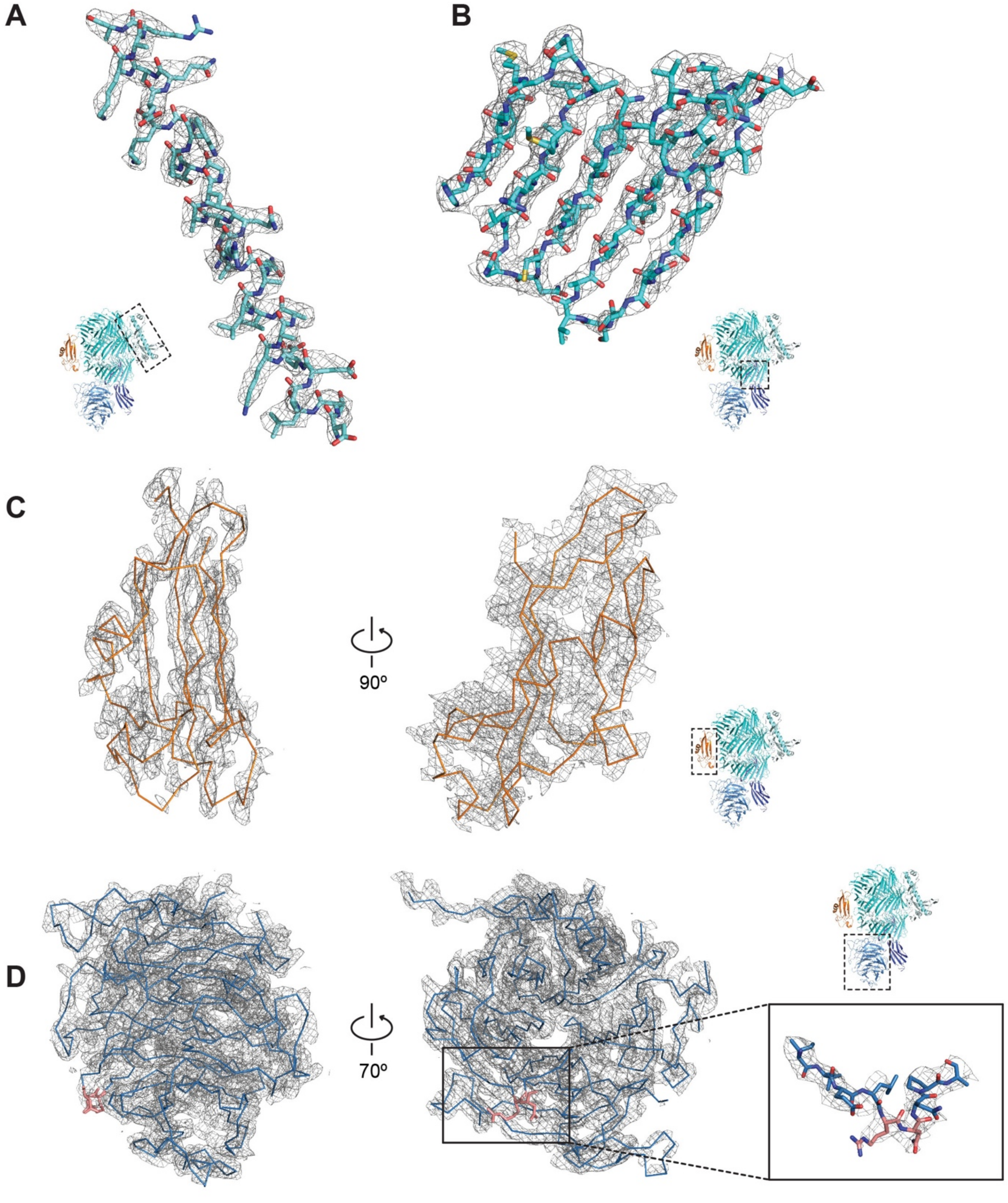


**Figure S4. Model quality, Related to Figure 1**

Snapshots of map versus model agreement. Each region is highlighted by a dashed box including the representative regions of an $\alpha$-helix in Tenm2 Tox-like domain (A), a $\beta$-sheet in Tenm2 $\beta$-barrel domain (B), the Lec domain of Lphn3 (C), and the entire Tenm2 $\beta$-propeller domain (D). The alternative splice site in $\beta$-propeller domain is located between Arginine 1156 and Asparagine 1157 (highlighted in pink). Contour level for individual snapshots: A, 5.0 rmsd; B, 5.0 rmsd; C, 4.0 rmsd; D, 5.0 rmsd.

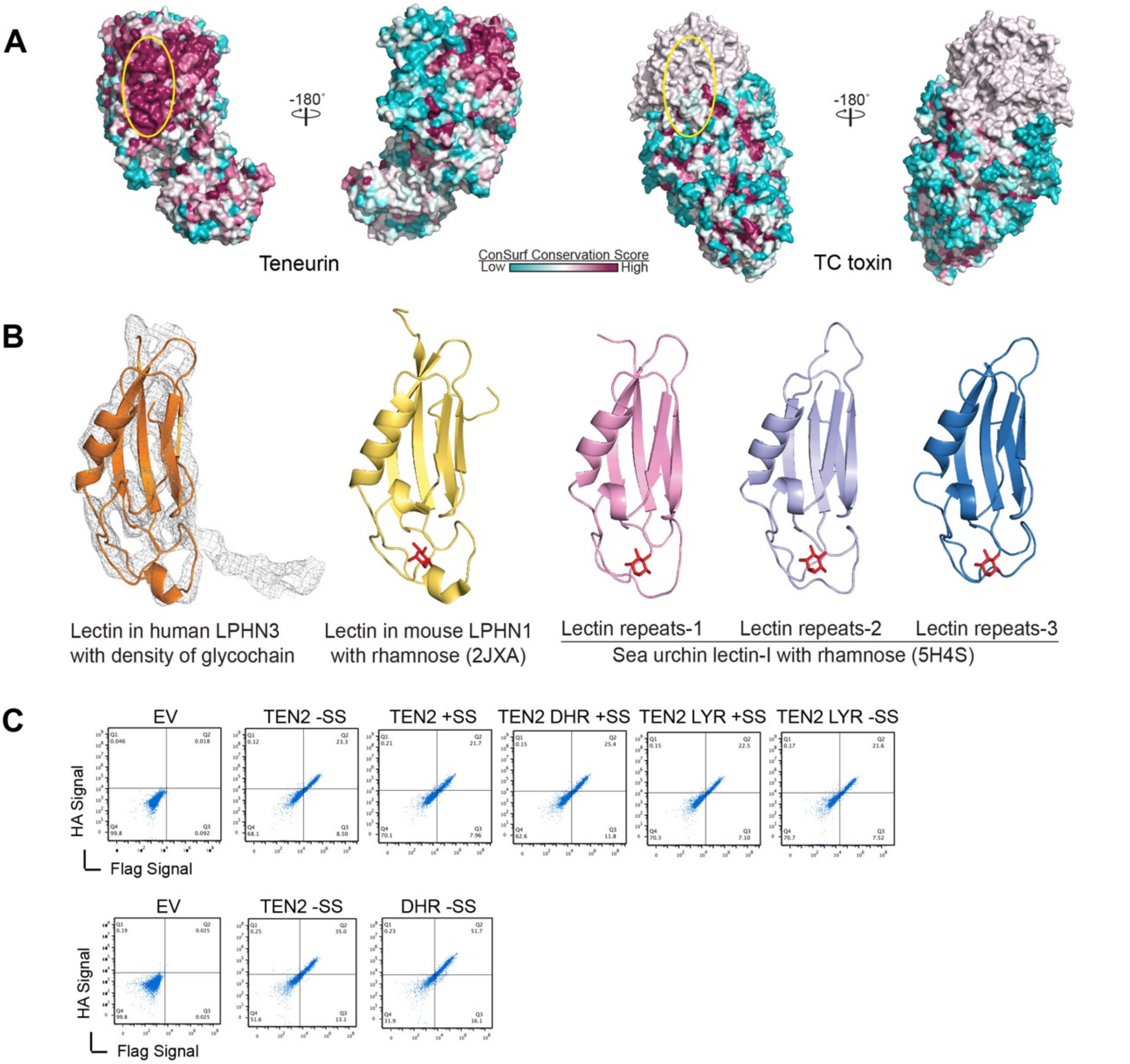


**Figure S5. Conservation, glycosylation and mutagenesis of the Lphn-binding site on Tenm2, Related to Figure 3**

(A) Conservation analysis of Tenm2 shows a highly conserved patch on the barrel surface (indicated by yellow circles) opposite to the Tox-like domain. This conserved patch is not present in bacterial toxins, implying the evolutionary importance of this region in Tenm2, which may be employed to interact with partner proteins. (B) Comparison of potential sugar binding site of Lphn3 Lec domain with other SUEL-related Lec family members. Lec domain in human Lphn3 with density of glycochain from human Tenm2 shown in grey mesh is colored as orange; Lec domain in mouse Lphn1 from NMR model (PDB: 2JXA) with rhamnose highlighted in red is colored as yellow (Vakonakis et al., 2008). Three Lec repeats in sea urchin Lphn3 (PDB: 5H4S) with rhamnose highlighted in red are colored as pink, palecyan and blue, respectively (Hatakeyama et al., 2017). All the Lec domains are aligned. Note that the glycochain density from Tenm2 is close to the sugar binding pocket of other Lec domains. (C) Flow cytometry analysis of cell-surface expression for full length Tenm2 on non-permeabilized HEK293T mammalian cells, compared to empty vector-transfected cells (EV) and cells that were transfected with mutant Tenm2 constructs.

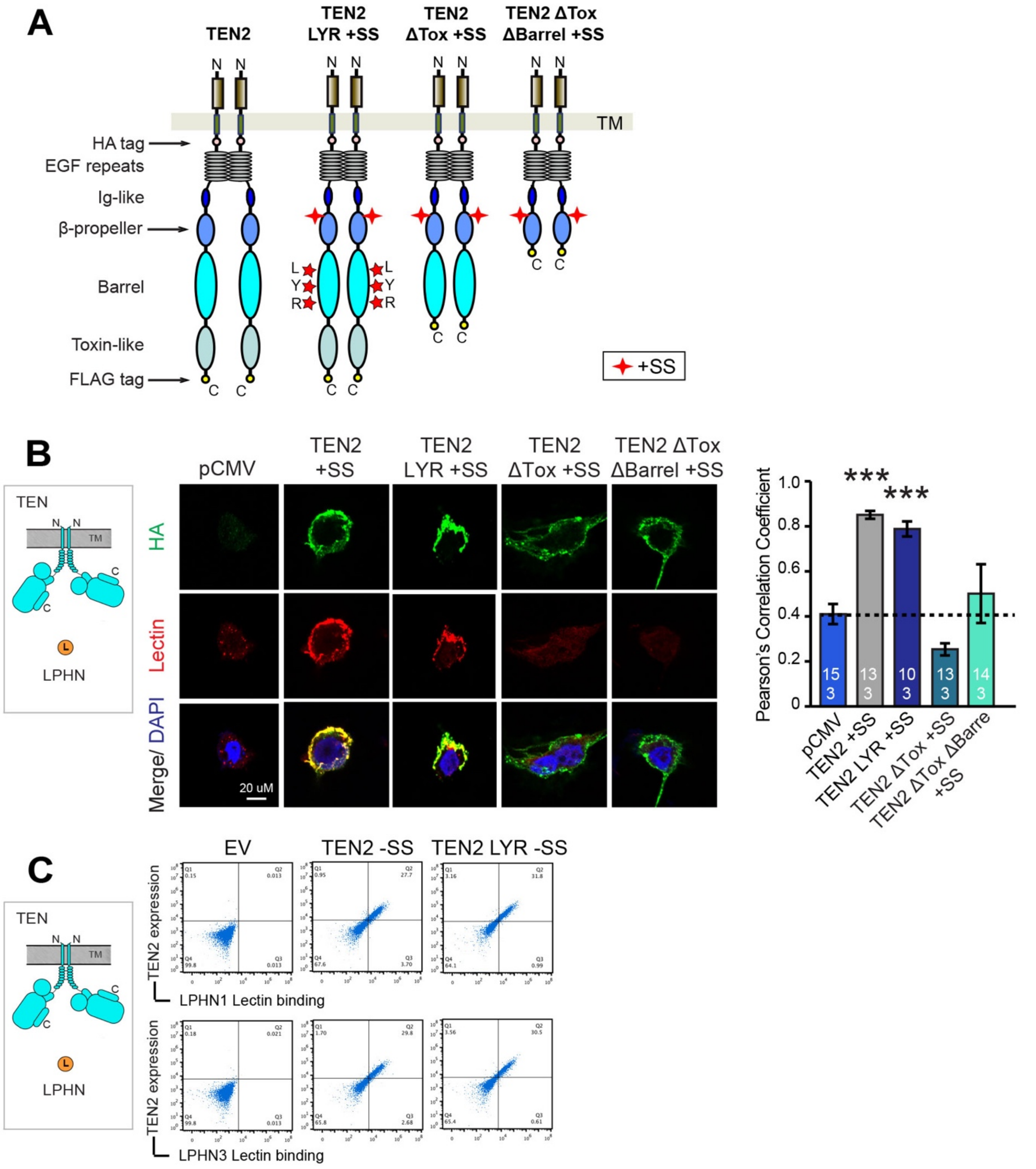


**Figure S6. LR mutation on Tenm2 has no influence on Lphn3 binding; Toxin-like domain deletion results, Related to Figure 3**

(A) Diagram for WT Tenm2 +SS, Tenm2 LR +SS (L1990N, R1992T), Tenm2 +SS Tox domain deletion (ΔTox) and Tenm2 +SS barrel and Tox domain deletion (ΔTox ΔBarrel) constructs that were used in this and our previous study (Li et al., 2018). Tenm2 LR +SS construct is generated as a positive control and it carries two

point mutations in the Tenm2 barrel which are not at the lphn-binding site. The Tenm2 +SS ΔTox and Tenm2 +SS ΔTox ΔBarrel are the same constructs as was used in Li et al.
(B) Cell surface staining assays suggest that the "LR" mutations on Tenm2 β-barrel has no influence on the binding between Tenm2 and Lphn3. Cell surface staining assays suggest that the toxin-domain deletion abolishes Lphn binding.
**Supplementary Discussion**: In Li et al., we reported that the toxin-like domain is needed for Lphn3 binding because the Tenm2 +SS ΔTox that deletes the toxin domain of Tenm2 had abolished Lec binding in cell-aggregation and cell-surface staining experiments and lacked any defects in cell-surface localization. These results mislead us to pinpoint the exact binding interface for Lec. We have repeated the experiments and confirmed these results are reproducible. We believe that this mutant escapes the protein quality control system and is localized on the cell surface although it is misfolded and thus is unable to bind Lphn. However, the real Lphn-binding site is not the toxin-like domain.
Scale bar indicates 20 um. Quantification of cell surface binding assays are shown next to the images.
(C) Wild-type (WT) and mutant Tenm2 proteins were tested for surface expression in HEK293T cells as well as their ability to bind soluble Lphn1/Lphn3 Lec domain using flow cytometry. Data relate to Tenm2 +SS delete Tox domain (ΔTox) and Tenm2 +SS delete Barrel and Tox domain (ΔTox Δ Barrel) are modified from Li et al., 2018 and separated by black line.

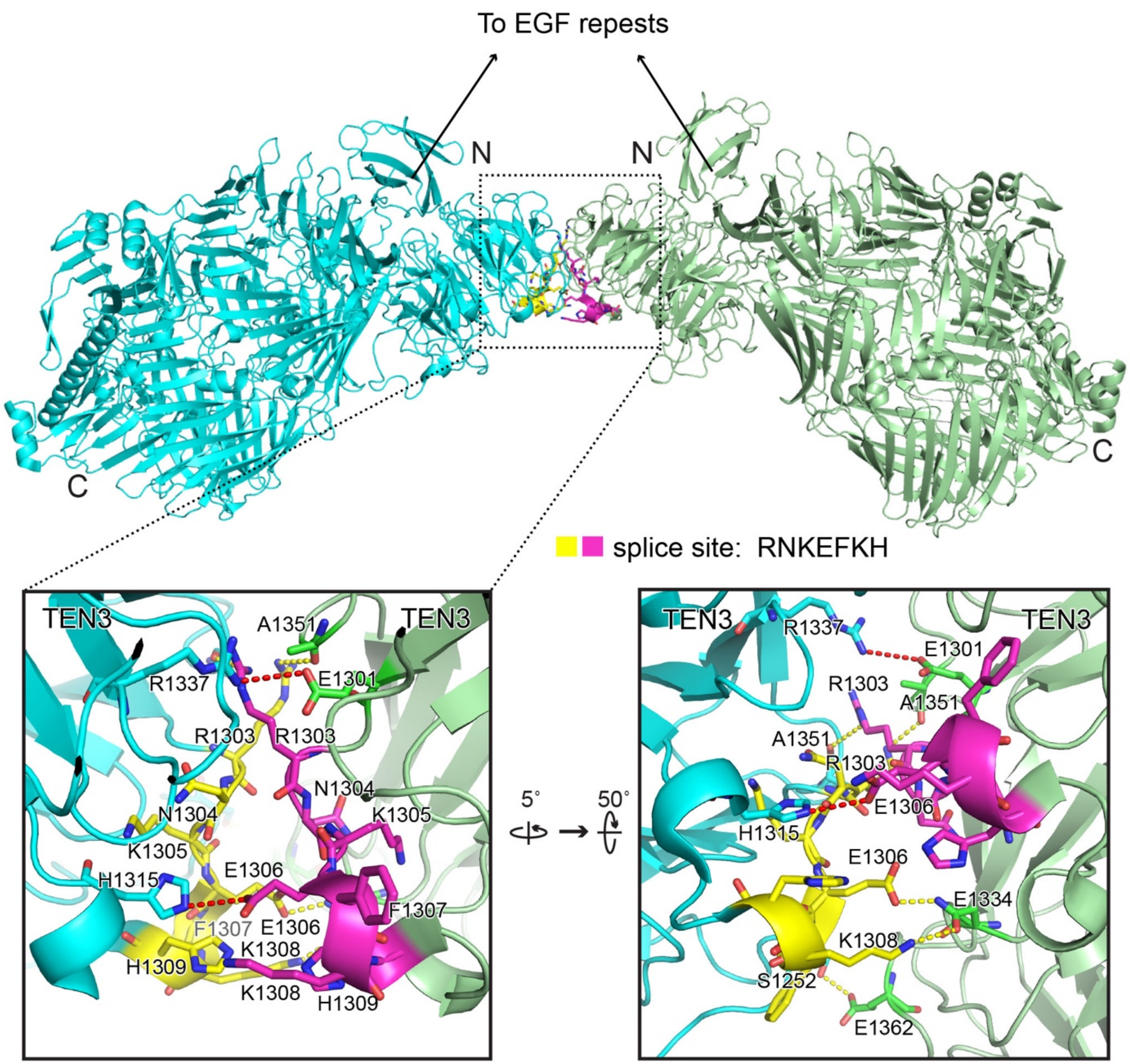


**Figure S7. Alternatively-spliced insert within the β-propeller mediates the Tenm2 +SS dimer interface, Related to Figure 7**

Structure of Tenm2 +SS dimer shows the splice inserts from each protomer (yellow and magenta residues) creates a binding interface and leads to Tenm2 dimerization via the beta-propeller. Close-up views of the dimer interface show two salt bridges and five hydrogen bonds are at the interface. One of the salt bridges is directly mediated by the glutamate (E1306) within the splice insert NKEFKHS and four of the hydrogen bonds also require splice site residues. Two salt bridges align almost parallel to each other and to the disulfide bonds between the EGF repeats and restricts the conformational flexibility of the Tenm2 head significantly (see Figure 7). The N-termini of both protomers face the same direction towards the EGF repeats, and thus, the dimer is positioned as a cis-dimer that will extend the zippering of the already existing EGF-mediated cis-dimer, although it was reported to form as a trans-homodimer, previously (Jackson et al., 2018). TEN protomers (PDB: 6FB3) are colored as cyan and palegreen, respectively, and splice sites are colored as tellow and magenta, respectively.
